# In-depth characterization of the serum antibody epitope repertoire in inflammatory bowel disease using phage-displayed immunoprecipitation sequencing

**DOI:** 10.1101/2021.12.07.471581

**Authors:** Arno R. Bourgonje, Sergio Andreu-Sánchez, Thomas Vogl, Shixian Hu, Arnau Vich Vila, Ranko Gacesa, Sigal Leviatan, Alexander Kurilshikov, Shelley Klompus, Iris N. Kalka, Hendrik M. van Dullemen, Adina Weinberger, Marijn C. Visschedijk, Eleonora A. M. Festen, Klaas Nico Faber, Cisca Wijmenga, Gerard Dijkstra, Eran Segal, Jingyuan Fu, Alexandra Zhernakova, Rinse K. Weersma

**Author notes:** Correspondence Prof. Rinse K. Weersma, M.D. Ph.D. Department of Gastroenterology and Hepatology University of Groningen and University Medical Center Groningen P.O. Box 30.001 9700 RB Groningen, the Netherlands. These authors contributed equally (Arno R. Bourgonje, Sergio Andreu-Sánchez). These authors contributed equally (Jingyuan Fu, Alexandra Zhernakova, Rinse K. Weersma).

## Abstract

Inflammatory bowel diseases (IBD), e.g. Crohn’s disease (CD) and ulcerative colitis (UC), are chronic immune-mediated inflammatory diseases. A comprehensive overview of an IBD-specific antibody epitope repertoire is, however, lacking. We leveraged a high-throughput phage-displayed immunoprecipitation sequencing (PhIP-seq) workflow to identify antibodies against 344,000 antimicrobial, immune and food antigens in 497 IBD patients as compared to 1,326 controls. IBD was characterized by 373 differentially abundant antibodies (202 overrepresented and 171 underrepresented), with 17% shared by both IBDs, 55% unique to CD and 28% unique to UC. Antibodies against bacterial flagellins dominated in CD and were associated with ileal involvement, fibrostenotic disease and anti-*Saccharomyces cerevisiae* antibody positivity, but not with fecal microbiome composition. Antibody epitope repertoires accurately discriminated CD from controls (AUC=0.89), and similar discrimination was achieved when using only ten antibodies (AUC=0.87). IBD patients thus show a distinct antibody repertoire against selected peptides, allowing patient stratification and discovery of immunological targets.

## Introduction

Inflammatory bowel diseases (IBDs), encompassing Crohn’s disease (CD) and ulcerative colitis (UC), are chronic inflammatory diseases of the gastrointestinal tract that are characterized by an inappropriate and uncontrolled immune response triggered by the gut microbiota in genetically susceptible individuals (Chang, 2020). A complex interplay between inherited and environmental factors, the gut microbiota and the host immune system is considered causative in the pathogenesis of IBD (Jostins et al., 2012; de Souza et al., 2017). The gut microbiota of patients with IBD show alterations in their composition and functionality, in particular a decrease in microbial diversity and the abundance of commensal, butyrate-producing bacteria and increased proportions of potentially pathogenic bacteria (Franzosa et al., 2019; Vich Vila et al., 2018; Frank et al., 2007). An immense traffic of luminal antigens occurs across the human intestinal epithelium, representing an interface between the mucosal immune system and the luminal microenvironment, where antigens are continuously sampled as an immune surveillance mechanism (von Martels et al., 2017). In the context of IBD, intestinal barrier function is compromised, resulting in excessive passage of luminal antigens (e.g. bacterial translocation) that elicits mucosal and systemic immune activation, which may in turn trigger and aggravate intestinal inflammation (Bischoff et al., 2014; Salim et al., 2011).

The human gut microbiota contains a variety of genes and species that represent a tremendous amount of potential antigens (Li et al., 2020; Pasolli et al., 2019). However, it is challenging to fully capture the complete set of antigens recognized by the human antibody repertoire because conventional technologies (e.g. enzyme-linked immunosorbent assays [ELISAs] or peptide arrays) do not allow for such large-scale measurements and are usually limited to parallel measurements of only a few hundreds to thousands of antigens. Moreover, although B-cell receptor sequencing (BCR-seq) methods report on the clonal diversity of the BCR DNA sequences that underlie antibody specificity, the exact nature of the antigens bound and their associations with IBD remain incompletely addressed (Vogl et al., 2021; Sterlin et al., 2020; Fadlallah et al., 2019). Phage-displayed immunoprecipitation sequencing (PhIP-seq) is a high-throughput technology that now allows for systematic investigation of hundreds of thousands of antigens simultaneously, and it has been successfully applied in both healthy populations and in the context of viral infections, although there is little data available on immune-mediated diseases (Vogl et al., 2021; Mohan et al, 2018; Zeevi et al., 2015; Larman et al., 2013; Mina et al., 2019; Xu et al., 2015). PhIP-seq is based on antigen libraries encoded by synthetic DNA oligonucleotides that are displayed on bacteriophages. These phages are incubated with human blood to initiate antigen–antibody pairing with the reactive antibodies present in the sample, and these antibodies are subsequently extracted by immunoprecipitation and sequenced for identification, resulting in an in-depth characterization of the blood antibody epitope profile (Vogt et al., 2012; Mohan et al., 2018).

In this study, we aimed to characterize the serum antibody epitope repertoire in patients with IBD *versus* healthy controls representing the general population and to associate these antibody epitope repertoires to relevant patient phenotypes, including clinical data and IBD-specific parameters, and to their fecal metagenomes. Using PhIP-seq technology, we established specific immune-based biomarker signatures in patients with IBD and identified distinct profiles for specific disease phenotypes.

## Results

### Cohort description

Demographic and clinical characteristics (IBD: *n* = 497; population controls: *n* = 1,326) are presented in Table 1. In total, 497 patients were included: 256 diagnosed with CD, 207 with UC and the remaining 34 patients with an undetermined type of IBD (IBD-U). A schematic overview of the study workflow is presented in Figure 1.

**Figure 1.**
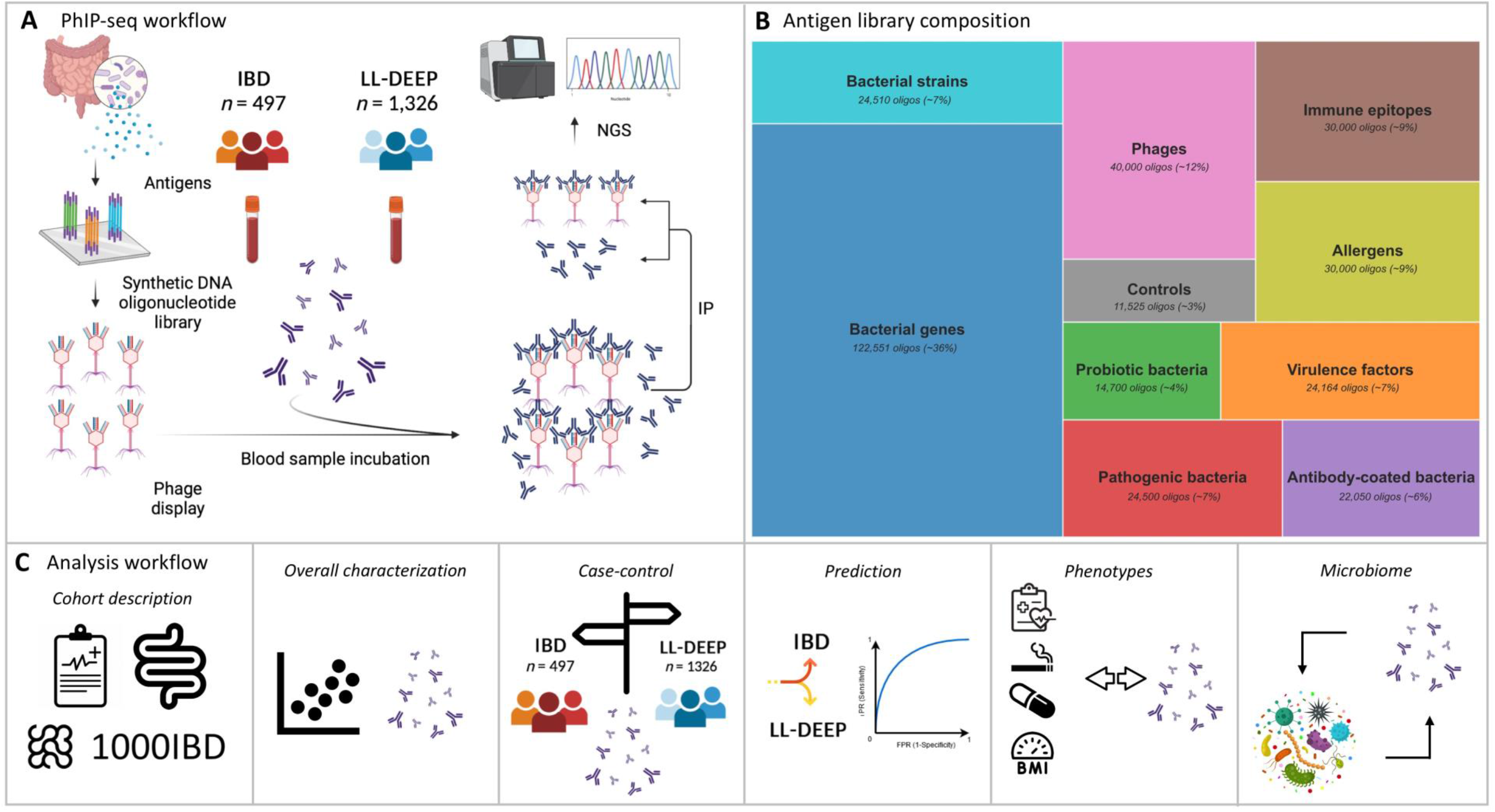
Methodological workflow of the study. (**A**) Antigens derived from bacterial genes and species, pathogenic, antibody-coated and probiotic bacteria, virulence factors, phages, allergens, immune epitopes and antigens for experimental controls (covering a total of 344,000 antigen peptides) were integrated into a synthetic DNA oligonucleotide library and displayed on bacteriophages. After addition of human blood samples, reactive antibodies from these samples bind to their corresponding antigens present in the library. After incubation, antibody-bound phages are extracted and separated from the unbound antibodies following immunoprecipitation (IP). Finally, antibody-bound phages are amplified in a high-throughput manner using next-generation sequencing (NGS). (**B**) Tree map visualization showing the composition the microbiota antigen phage library. Approximately 36% of the full library consisted of antigens of bacterial genes (derived from metagenomics sequencing data), followed by phages, allergens and immune epitopes (from the Immune Epitope Database, IEDB), bacterial strains, pathogenic bacteria, virulence factors (from the virulence factor database, VFDB), antibody-coated bacterial species, probiotic bacteria and several biological and technical controls (see *Supplementary Methods* for details). (**C**) Analysis of the PhIP-seq data was performed in steps. First, a cohort description was provided consisting of demographic and clinical characteristics of patients with IBD. Subsequently, an overall characterization of the data was generated by calculating summary statistics of the antibody epitope repertoires and performing principal component analysis (PCA). This step was followed by case–control analyses in which significantly bound peptides were compared between patients with IBD and healthy individuals while controlling for the effect of age and sex. Predictions of the diagnosis of CD or UC using the antibody epitope repertoires were established by evaluating classification accuracy. Antibody epitope repertoires were then analyzed in relation to (IBD-specific) phenotypes, e.g. Montreal disease classification and surgical history. Finally, the concordance between blood antibody epitope repertoires and fecal microbiome data (metagenomics sequencing) was assessed. Abbreviations: GWAS, genome-wide association study; HLA, human leukocyte antigen; IBD, inflammatory bowel disease; IP, immunoprecipitation; LL-DEEP, Lifelines-DEEP cohort; NGS, next-generation sequencing.

**Table 1.**
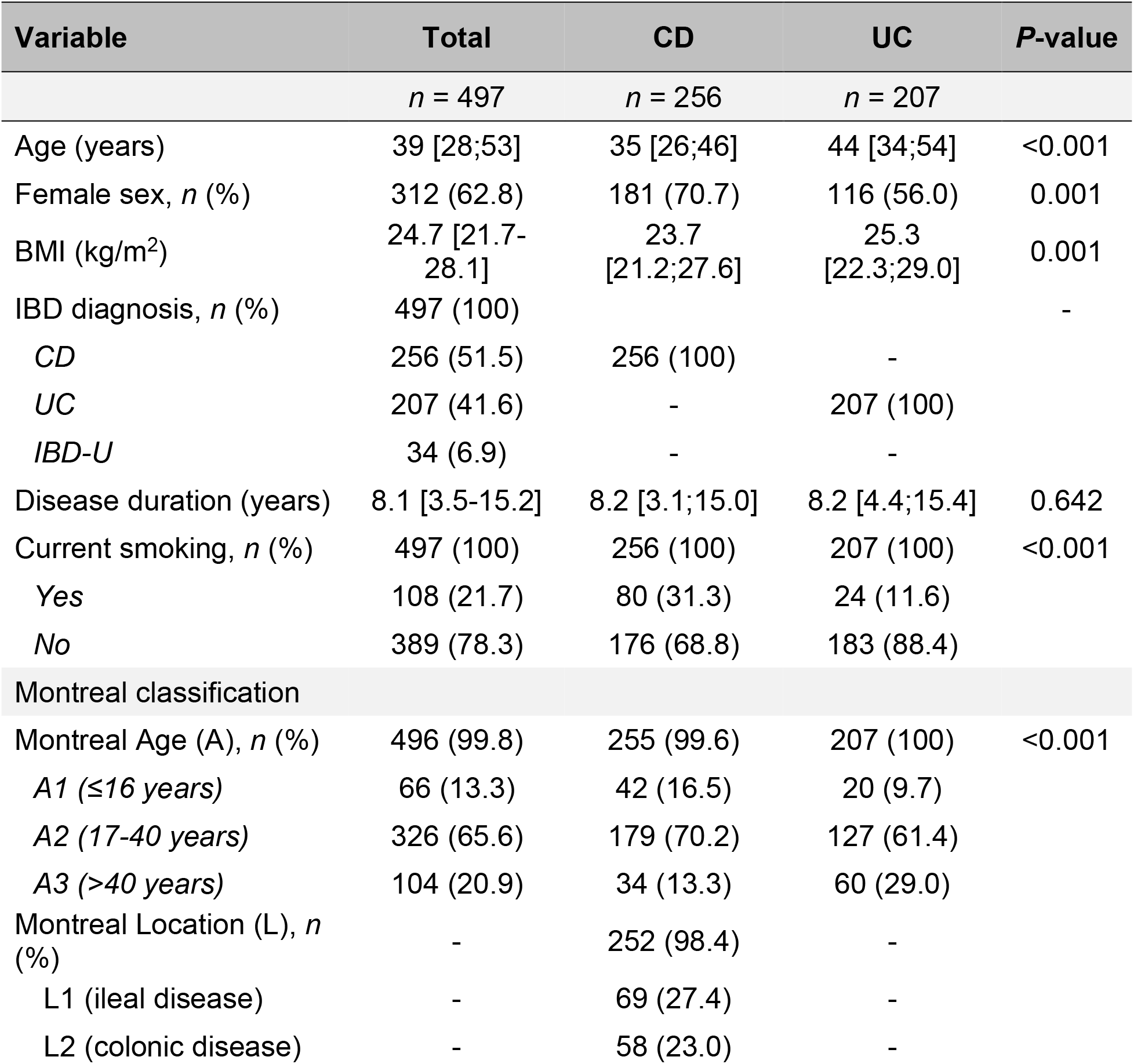
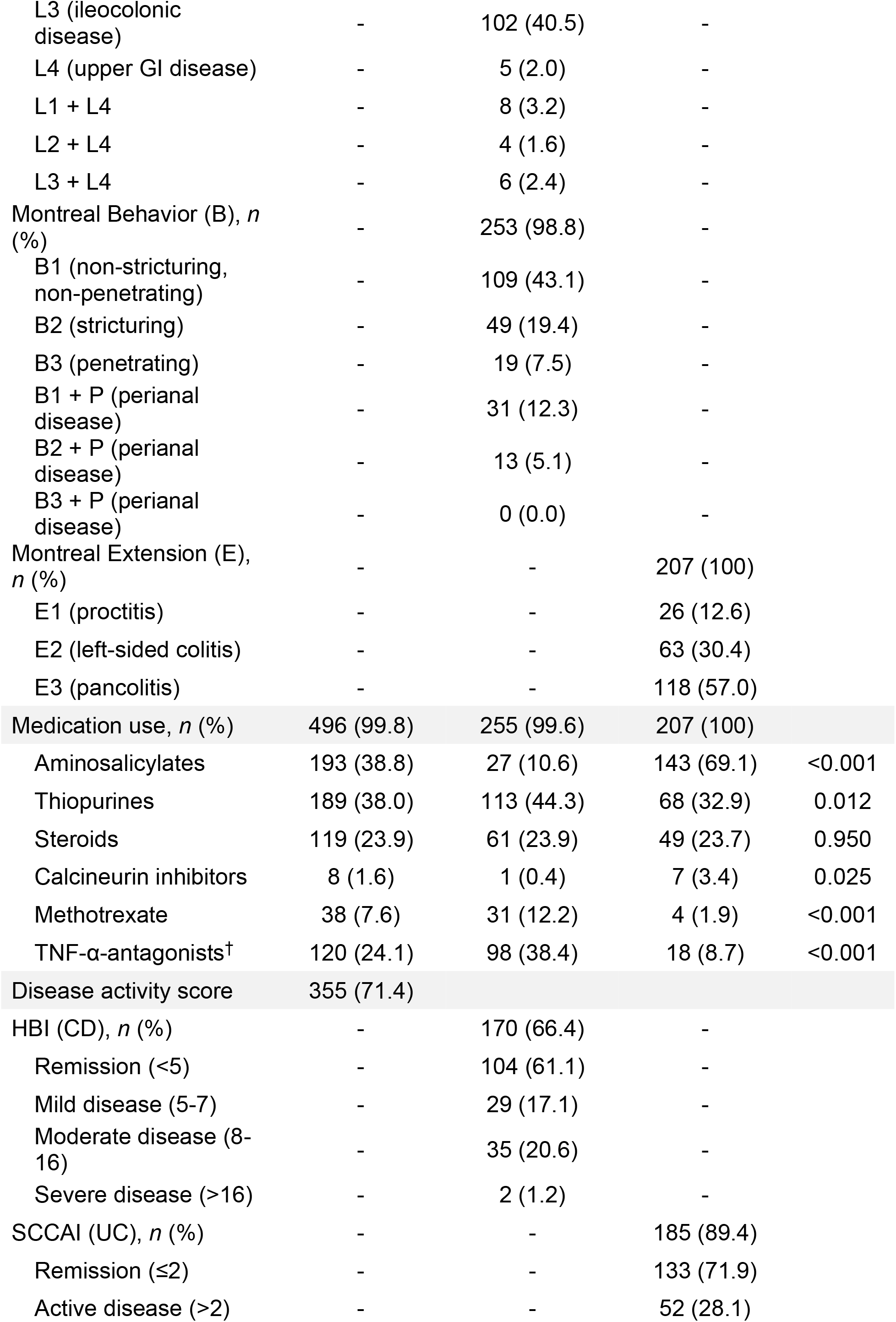
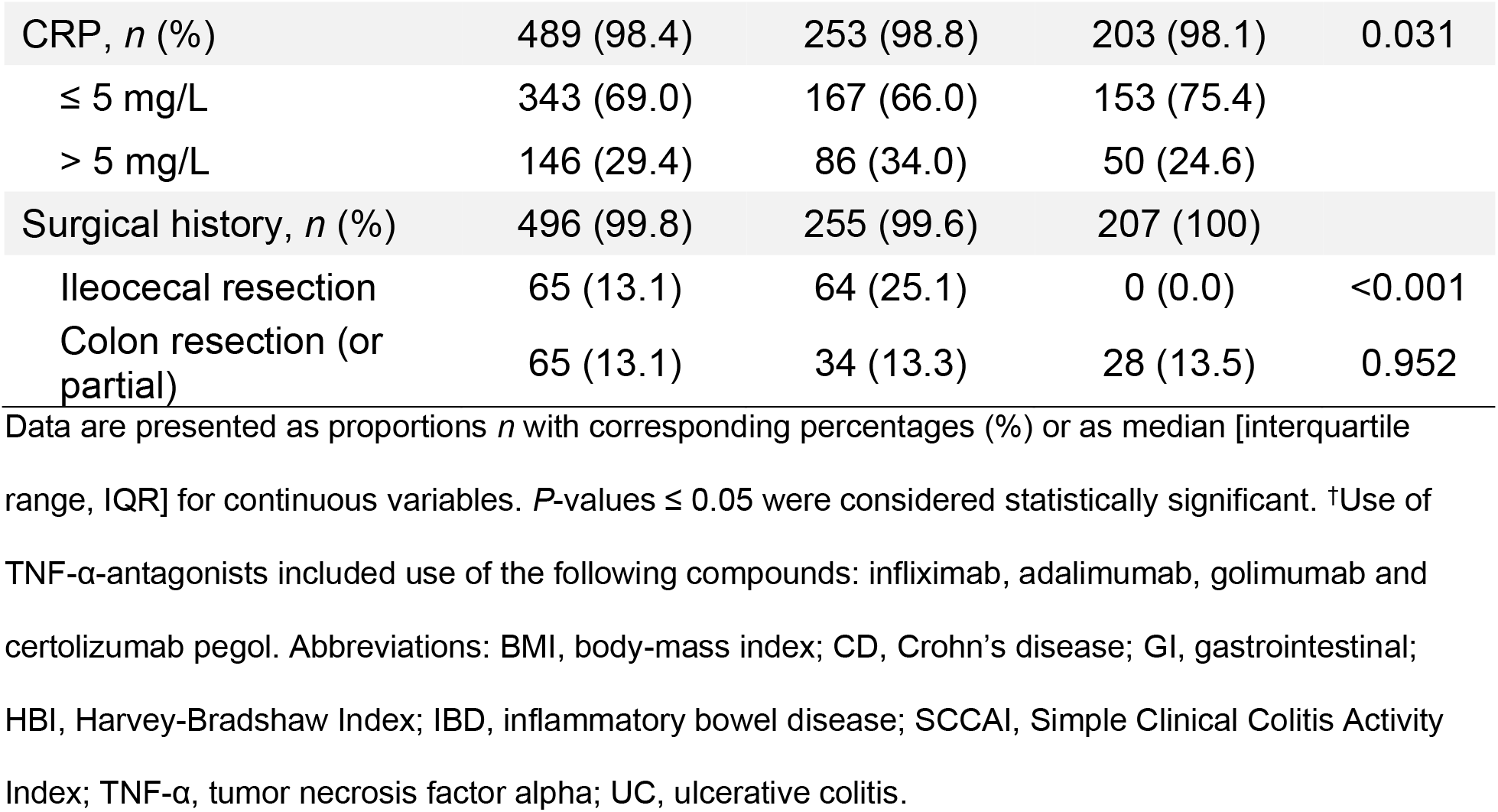
Demographic and clinical characteristics of the study population (IBD: *n* = 497).

### Overall characterization of serum antibody epitope repertoires in patients with IBD

We performed dimensionality reduction analyses (principal component analysis, PCA) to visualize the heterogeneity of the antibody epitope repertoires in patients with IBD (Figure 2). Two distinct clusters were observed based on the first two principal components (PCs) (Fig. 2A-C), and these were further confirmed by a *k*-means clustering analysis (Fig. 2C). Antibody epitope diversity (i.e. the number of enriched antibody-bound peptides per patient) clearly explained the variation in both PC1 and PC2 (both *P* < 0.001). After comparing antibody-bound peptides between both cluster identities, this clustering was found to be mainly driven by antigens derived from cytomegalovirus (CMV). This observation was confirmed when comparing both clusters for matching serological CMV measurements (measured by ELISA for *n* = 297 patients, *R* = 0.91, *P* < 0.001, Suppl. Fig. S1A) and was also in line with the clustering observed in a population-based cohort (Andreu-Sánchez et al., manuscript in preparation). No significant batch effects were observed, as the IDs of plates onto which samples were loaded were not associated to the first two PCs or the subsequent eight PCs, which cover a cumulative explained variance of 21% in the antibody data (Suppl. Fig. S1B). The median number of positive antibody-bound peptides per individual was 1,015 (IQR: 721–1362, full range: 179–2722). Overall antibody epitope diversity was similar for patients with CD and UC (median with IQR: 1015 [721;1365] and 1015 [721;1341], respectively, *P* = 0.325, Fig. 2D). More than 100,000 antibody-bound peptides were positive in at least one individual, whereas only a few peptides were shared among all patients (Fig. 2E). These observations, which indicate the existence of both individual-specific and population-wide antibody responses, are in line with recent studies that performed PhIP-seq analysis in population-based cohorts (Vogl et al., 2012; Andreu-Sánchez et al. manuscript in preparation). In fact, just 2,368 different antigen peptides were present in 5–95% of individual patients with IBD (Fig. 2F), with 83% of peptides shared between both IBD subtypes and the remaining 17% specific to either CD or UC.

**Figure 2.**
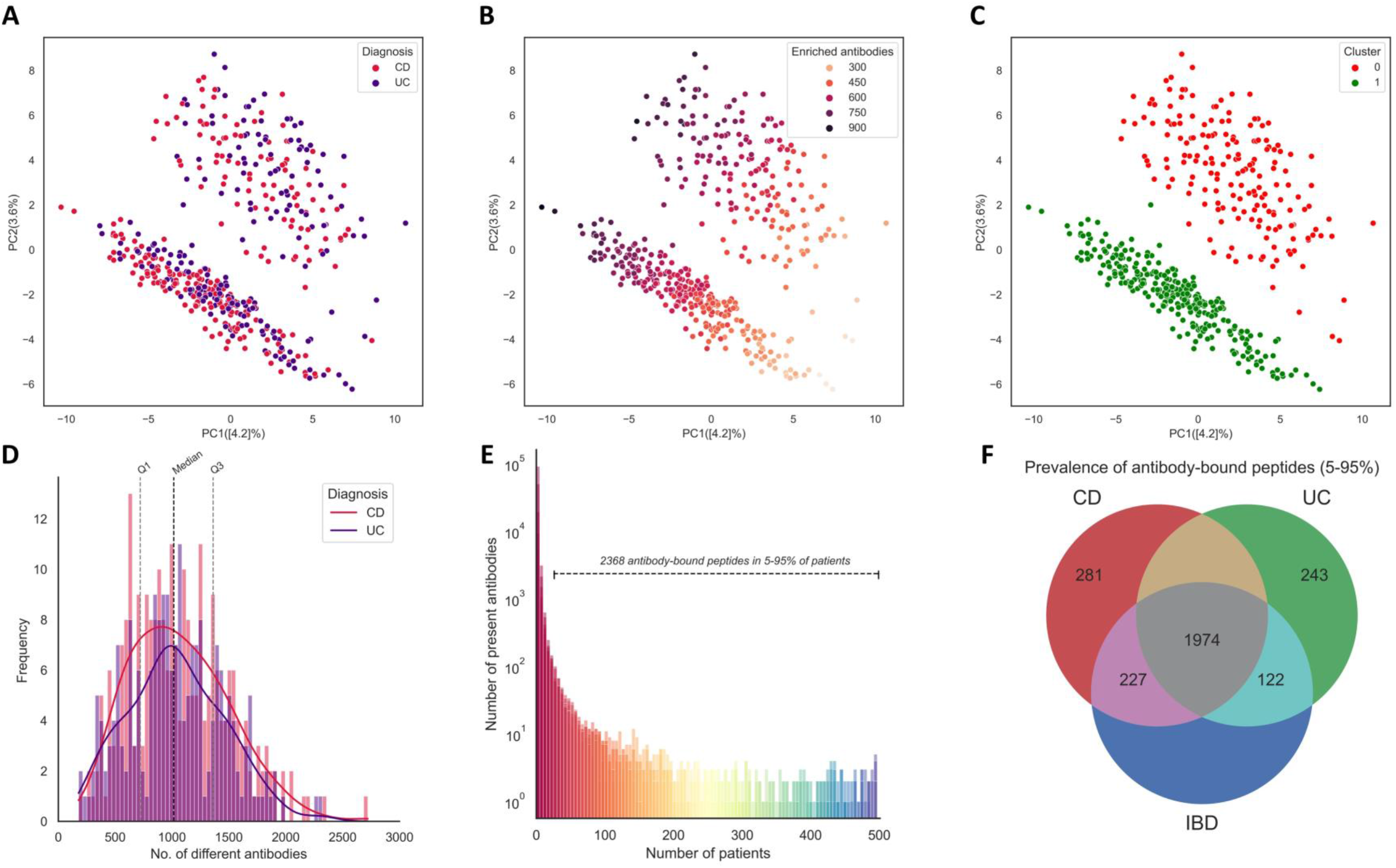
Overall characterization of serum antibody epitope repertoires in patients with IBD. (**A–C**) Principal component analysis (PCA) plots demonstrating the first two principal components (PCs) describing data variation. (**A**) IBD diagnosis could partially explain PC1, but was not associated to PC2. (**B**) Antibody diversity, i.e. the number of significantly enriched antigen peptides per patient, was significantly associated to both PCs. (**C**) A *k*-means clustering algorithm (*k* = 2) successfully separated two distinct clusters, and these appeared to be based on the enrichment of antigen peptides from herpes virus proteins, particularly cytomegalovirus. (**D**) Antigen peptide diversities among patients with CD and UC show comparable distributions, with a median of 1,015 peptides per patient. Lines indicate kernel density estimates of the CD-and UC-specific distributions. (**E**) Prevalence of antigen epitope repertoires amongst all patients, showing that tens of thousands of antigen peptides were found in only a single patient, while only few peptides were shared among the full study cohort. (**F**) Venn diagram showing the prevalence of significantly enriched antigen peptides amongst patients with CD (*n* = 256) and UC (*n* = 207) (diagnoses of IBD-U excluded here), indicating that a large proportion of peptides are shared among patients with CD and UC. Abbreviations: CD, Crohn’s disease; PC, principal component; Q1, first quartile (25^th^ percentile); Q3, third quartile (75^th^ percentile); UC, ulcerative colitis.

### Patient age and sex confirm known associations with antibody epitope repertoires

Next, we examined the associations between antibody responses and patient age and sex, as these factors have previously been shown to be strongly associated with antibody epitope repertoires (Vogl et al., 2012). We compared the oldest patients (Q4, >53 years old, *n* = 115) to the youngest patients (Q1, <28 years old, *n* = 115) and identified significant associations to 199 different antibody-bound peptides (false discovery rate (FDR) < 0.05) (**Table S1**). Older patients with IBD showed increased antibody responses against proteins from several herpesviruses (herpes simplex virus type 1 [HSV-1]/HHV-1, varicella-zoster virus [VZV]/HHV-3 and cytomegalovirus [CMV]/HHV-5), hepatitis A virus proteins and proteins from the order Bacteroidales. Younger patients with IBD showed increased antibody responses against proteins of Shiga-like toxin-producing types of *Escherichia coli* (e.g. serotype O157:H7), including the EspD secretor protein, translocon proteins and the translocated intimin receptor (Tir). Younger IBD patients also more frequently demonstrated antibodies against some flagellin-coated bacteria from the Clostridiales order, rhinoviruses (particularly serotype 2) and autolysins of *Staphylococcus aureus*. Sex was significantly associated with two antigen peptides (FDR < 0.05) (**Table S2**). Female patients with IBD more frequently exhibited antibody responses against antigens from *Lactobacillus acidophilus* bacteria, some of which were also among the top nominally significantly associated antigen peptides (*P* < 0.05), an association previously reported for a population-based cohort (Vogl et al., 2012). Male patients with IBD showed increased antibody responses against surface fibrils and adhesins of *Haemophilus influenzae* bacteria. Most of these associations between patient age and sex and antibody-bound peptides correspond very well with those reported in recent population-based cohort studies (Vogl et al., 2012; Andreu-Sánchez et al., manuscript in preparation).

### Distinct antibody responses in patients with IBD compared to healthy individuals

A comparison of the antibody epitope repertoires of patients with IBD and healthy controls (Andreu-Sánchez et al., manuscript in preparation), while controlling for the effects of age and gender, revealed 373 differentially abundant antibody-bound peptides, of which 202 were overrepresented and 171 were underrepresented in IBD compared to healthy controls (FDR < 0.05) (Fig. 3A, **Tables S3-5**). Patients with CD accounted for most differentially abundant peptides (55%) as compared to patients with UC (28%), while approximately one-fifth were shared between both IBD subtypes (17%). Results from these comparative analyses did not change substantially when an age-and sex-matched, equally sized subset of the healthy controls was selected for both patients with CD and UC (**Tables S6–7**). Patients with CD showed a distinct overrepresentation of antibody-bound peptides derived from bacterial flagellins (Fig. 3B, **Table S4**, **Table S8**). These consisted of significantly enriched flagellin antigen peptides from bacterial taxa belonging to the order Clostridiales and family Lachnospiraceae, including *Roseburia*, *Eubacterium*, *Clostridium* and *Agathobacter* spp., as well as to taxa belonging to presumed pathogenic bacteria, including those belonging to the families Legionellaceae (e.g. *Legionella pneumophila*), Borreliaceae (e.g. *Borrelia burgdorferi*) and Burkholderiaceae (e.g. *Burkholderia pseudomallei*). Apart from flagellin proteins, higher frequencies of antibody-bound peptides derived from viruses were found in patients with CD, including nuclear antigens of Epstein-Barr virus (EBV) and phosphoproteins of CMV. In addition, antibody-bound peptides representing other viruses (e.g. genome polyproteins of enterovirus type B and C, non-structural polyproteins of Norwalk virus), bacterial cell wall components (e.g. adhesins of *Haemophilus influenzae*, autolysins such as N-acetylmuramoyl-L-alanine amidase of *Staphylococcus aureus*, and surface proteins of *Streptococcus pneumoniae* and *Streptococcus pyogenes*), and several types of virulence factors (including autotransporter proteins of *H. influenzae*, *E. coli* and *Neisseria meningitides*, translocator proteins of *E. coli* and *Pseudomonas aeruginosa* and anti-phagocytic M proteins of *S. pyogenes*) occurred less frequent in patients with CD compared to healthy individuals. In addition, antibody responses against wheat allergens were found to be more frequent in patients with CD, including allergens from *Triticum aestivum*, (common wheat), *Secale cereale* (rye), *Zea mays* (maize corn), *Aegilops tauschii* (wheat), *Hordeum vulgare* (barley), and *Oryza sativa* (Asian rice). In contrast, egg allergens (ovomucoid peptides from chickens, *Gallus gallus*) were decreased in patients with CD compared to healthy individuals. Finally, few antibody responses against human proteins (i.e. autoantigens) were observed to be more frequent in CD, e.g. peptides from the alpha chain of collagen type IV.

**Figure 3.**
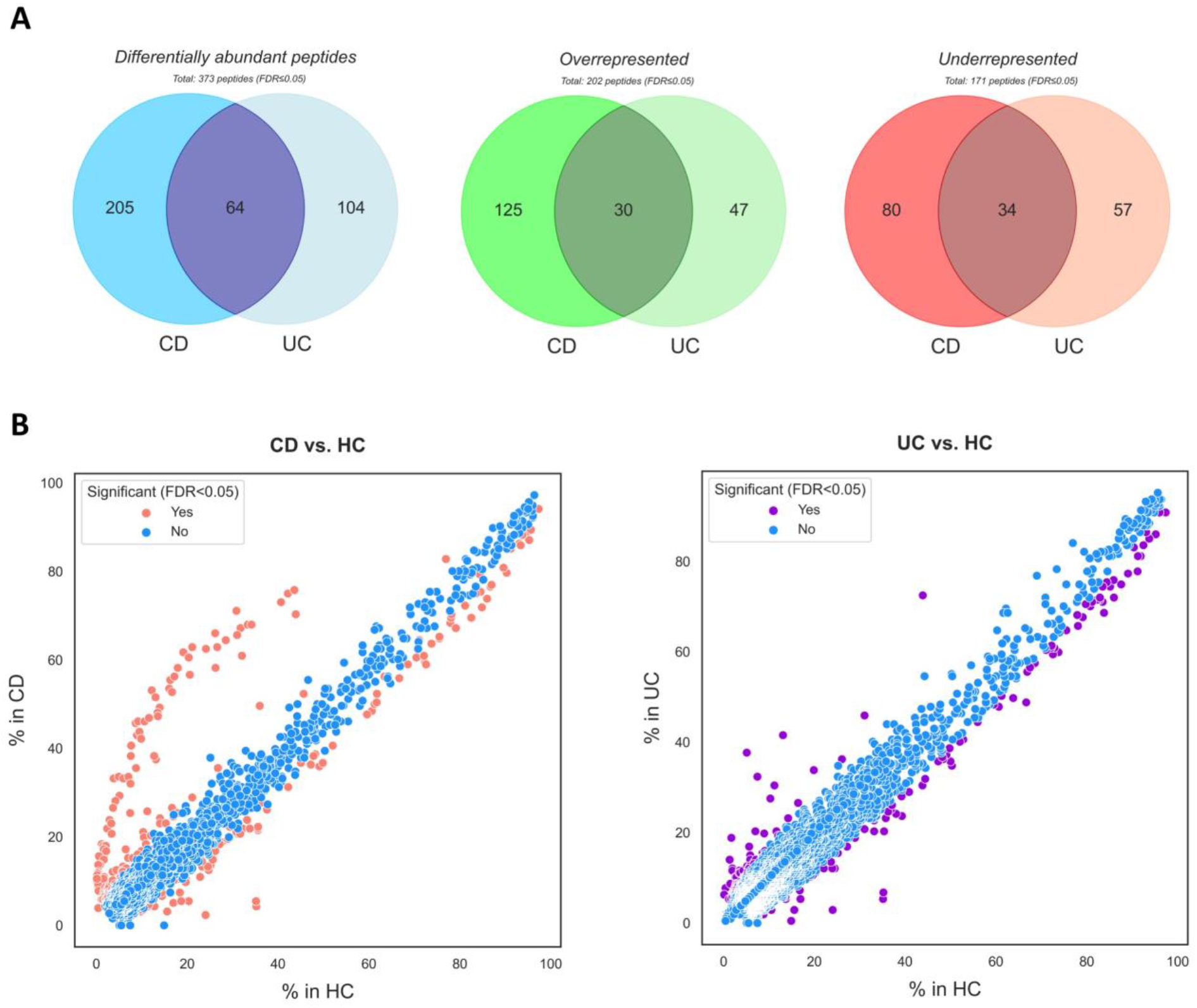
Distinct antibody epitope repertoires in patients with IBD compared to healthy individuals. (**A**) Case–control analysis of patients with IBD and healthy individuals, controlling for age, gender and plate ID, revealed 373 differentially abundant peptides (blue Venn diagram) in patients with IBD compared to healthy controls. Of these, 202 peptides were overrepresented in patients with IBD (green Venn diagram), whereas 171 peptides were underrepresented (red Venn diagram). (**B**) Differentially abundant antibody-bound peptides between patients with CD and UC compared to healthy controls (FDR < 0.05, logistic regression analysis, see **Tables S3–5** for full lists of differentially enriched peptides). Abbreviations: CD, Crohn’s disease; FDR, false discovery rate; HCs, healthy controls; UC, ulcerative colitis.

In contrast to CD, patients with UC did not demonstrate distinct anti-flagellin antibody responses but did exhibit an overrepresentation of antibody responses directed against fibronectin-binding proteins, including fibronectin-binding proteins A and B of *S. aureus* and fibronectin-binding proteins SfbII and A of *S. pyogenes*, as well as WxL domain-containing proteins (unknown function), OstA-like domain-containing proteins of *Parabacteroides johnsonii* (involved in lipopolysaccharide assembly), periplasmic beta-glucosidase BgIX of *E. coli* (involved in bacterial carbohydrate metabolism) and STN domain-containing proteins (involved in TonB-dependent active uptake of nutrients) of *Bacteroides dorei* (**Table S5**, Fig. 3B). However, some of these results warrant cautious interpretation, as coagulation-associated antibody-bound peptides may be differentially abundant due to the comparison of serum (which is generally depleted of coagulation proteins) *versus* plasma (which may still contain coagulation factors). For example, staphylococcal coagulase proteins were markedly underrepresented in UC compared to healthy controls (^∼^5% vs. ^∼^35%, respectively) (**Table S5**). To address this issue, we also compared antibody responses between patients with CD and UC using serum samples from both groups and observed similar results (**Table S8**). Patients with UC also demonstrated decreased antibody responses against pneumococcal histidine triad proteins and choline-binding proteins of *S. pneumoniae*, murein hydrolase activator proteins of *E. coli* (involved in peptidoglycan turnover, regulating cell wall growth), M proteins of *S. pyogenes* and several types of bacterial cell wall components (e.g. N-acetylmuramoyl-L-alanine amidase, adhesins of *H. influenzae* and other surface proteins). In addition, more frequent antibody responses against several viruses (among others, influenza A, hepatitis E, EBV, CMV, and rubella) were observed in patients with UC, concomitantly with decreased antibody responses against respiratory syncytial virus (RSV), rhinoviruses, and Norwalk viruses. Finally, patients with UC demonstrated increased antibody responses against a couple of human autoantigens, including the MAP kinase-activating death domain protein (apoptosis protein), aldolase B (carbohydrate metabolism enzyme), and an ATPase phospholipid transport protein, while they showed decreased antibody responses against the huntingtin protein (involved in axonal transport).

### Antibody epitope repertoires accurately discriminate between CD and healthy controls

Subsequently, we aimed to examine the discriminative capacity of antibody epitope repertoires with regard to the presence of CD *vs.* healthy controls, the presence of UC *vs.* healthy controls and the presence of CD *vs.* UC. A selection of 2,687 antibody-bound peptides (after excluding coagulation-associated peptides, *n* = 128) was used for classification between CD (*n* = 256), UC (*n* = 207) and equally sized age-and sex-matched subsets of healthy controls (Andreu-Sánchez et al., manuscript in preparation). A logistic regression model with elastic net penalty was fitted based on the antibody epitope repertoires using 80% of the data as training set, with the goal of classifying patients with CD or UC from healthy controls and patients with CD from patients with UC (Fig. 4, **Table S9**). When evaluating model performance on 20% of the data (test set), antibody epitope repertoires demonstrated a highly accurate discrimination between patients with CD and healthy controls (area under the curve (AUC) = 0.89, F1-score = 0.80) (Fig. 4A–B). In this classification, antibody epitope repertoires had a sensitivity of 76% and specificity of 84% in predicting the presence of CD at a default probability threshold of 0.5 (Fig. 4B), with a positive predictive value (PPV) of 83% and negative predictive value (NPV) of 78%. In contrast to CD, serum antibody epitope repertoires showed less, but still accurate, discrimination between patients with UC and healthy controls (AUC = 0.80, F1-score = 0.70). Here, sensitivity and specificity for the detection of UC were 71% and 68%, respectively, with a PPV and NPV of 69% and 70%, respectively. Finally, we assessed the predictive performance of antibody epitope repertoires between patients with CD and UC, which showed only moderate discriminative capacity (AUC = 0.68, F1-score = 0.66). Classification accuracies of all three discriminations were comparable across different machine learning methods (gradient boosting machine (GBM), support vector machine (SVM) and avNNet models), indicating that the results we observed were not specific to the methodology chosen (logistic regression with elastic net penalty) (**Table S9**). The relative importance of the antibody-bound peptides contributing to each of the presented classifications following elastic net regression can be found in **Table S10**.

**Figure 4.**
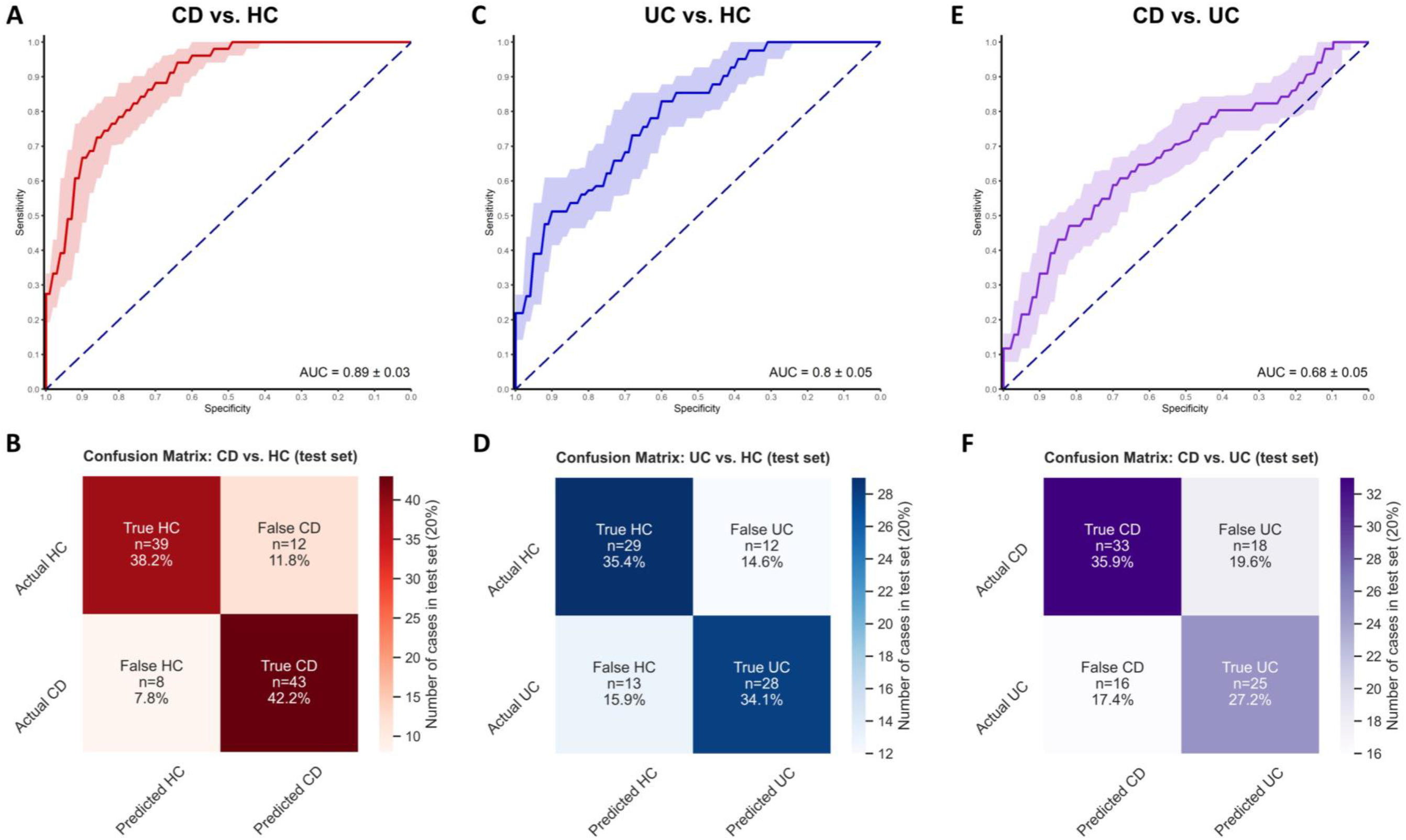
Classification between patients with CD and healthy controls, patients with UC and healthy controls and patients with CD and UC based on antibody epitope repertoires. Antibody epitope repertoires show superior discrimination between patients with CD and HC (**A, B**) in comparison to patients with UC vs. HC (**C, D**) or between patients with CD and UC (**E, F**). (**A,C,E**) ROC curves demonstrating the discriminative capacity of antibody epitope repertoires in classifying between patients with CD and HC in the test set (20% of the data). (**B,D,F**) Confusion matrices showing the predicted class numbers (from left to right and top to bottom: true negatives [TN], false positives [FP], false negatives [FN] and true positives [TP]) and proportions in the test set while adopting a probability threshold of 0.5. Abbreviations: AUC, area under the curve; CD, Crohn’s disease; HC, (age-and sex-matched) healthy controls; UC, ulcerative colitis.

### Patients with CD can be accurately identified based on only ten antibody-bound peptides

As a next step, we aimed to identify the top contributing antibody-bound peptides with regard to the three classification tasks (CD *vs*. healthy controls, UC *vs*. healthy controls and CD *vs*. UC) and evaluate the extent to which only a selection of antibody-bound peptides could discriminate between groups (i.e. without requiring a large number of antibodies). To do so, we adopted a feature selection method using a recursive feature elimination (RFE) procedure that fits a model and optimizes it by removing the weakest features (here: individual antibody-bound peptides) until the pre-specified number of antibody-bound peptides is reached (**Table S11–12**). When selecting the top-five antibody-bound peptides, an accurate discrimination between patients with CD and HCs could already be achieved (AUC = 0.81, F1-score = 0.74) (Figure 5A–B). When selecting the top-ten antibody-bound peptides, this discrimination improved considerably further (AUC = 0.87, F1-score = 0.77) and was statistically significant (DeLong’s test, Z-statistic: −3.25; FDR = 0.01) (**Table S13**). Notably, this classification model achieved similar discriminative performance to the model that included all contributing antibody-bound peptides (Fig. 4A), both of which indeed showed no statistically significant discriminative accuracies (DeLong’s test; Z-statistic 2.36; FDR = 1.00) (**Table S13**). The top-ranked antibody-bound peptides were the P30 adhesin protein of *Mycoplasma pneumoniae* (less frequent in CD), the human collagen type IV alpha chain protein, tegument protein of HSV-2, a translocator protein of *Pseudomonadacea*e, and flagellins from *Clostridiales*, *Legionellaceae*, *Eubacterium*, *Roseburia* and *Borrelia* (all more frequent in CD). When discriminating between patients with UC and healthy controls, a selection of the top-five and top-ten antibody-bound peptides already resulted in a reasonably accurate discrimination that approached that observed in the test set using the full set of antibody-bound peptides (top five: AUC = 0.75, F1-score = 0.66; top ten: AUC = 0.76, F1-score = 0.67) (Fig. 5C–D). The top contributing antibody-bound peptides to this classification were the P30 adhesin protein of *Mycoplasma pneumoniae* and tegument protein of HSV-2 (both overlapping with previous model, both less frequent in UC and CD); peptides belonging to surface proteins or zinc metalloprotease of *Streptococcus pneumoniae* and the EspB protein of *E. coli* O127:H6 (all less frequent in UC), WxL domain-containing protein of *Enterococcus faecalis*, DUF4988 domain-containing protein of *Bacteroides* and two allergen peptides belonging to *Danio rerio* (zebrafish) and *Bos* (wild or domestic cattle) (all more frequent in UC). Finally, we extracted the top five and top ten antibody-bound peptides contributing to the classification of patients with CD from patients with UC, which showed only moderate discriminative capacity (top five: AUC = 0.73, F1-score = 0.71; top ten: AUC = 0.69, F1-score = 0.67) (Fig. 5E–F), similar to that observed when using all entries of antibody-bound peptides. Among the top contributing antibody-bound peptides here were an autolysin of *Lactobacillus salivarius* (N-acetylmuramoyl-L-alanine amidase), a plant fungal allergen (*Alternaria alternata*) and flagellins from *Eubacterium* and *Legionellaceae* and the P200 protein of *Mycoplasma pneumoniae* (all more frequent in CD) and a *Bacillus* phage peptide and viral peptides including Norwalk virus, EBV and enterovirus B (all more frequent in UC). Apart from the individual antibody-bound peptides among these top selections, these classification analyses confirm that antibody-bound peptides from flagellin-coated bacterial species dominated among the antibody-bound peptides contributing to the classification of patients with CD from healthy individuals and from patients with UC (**Table S10**, **Table S12**).

**Figure 5.**
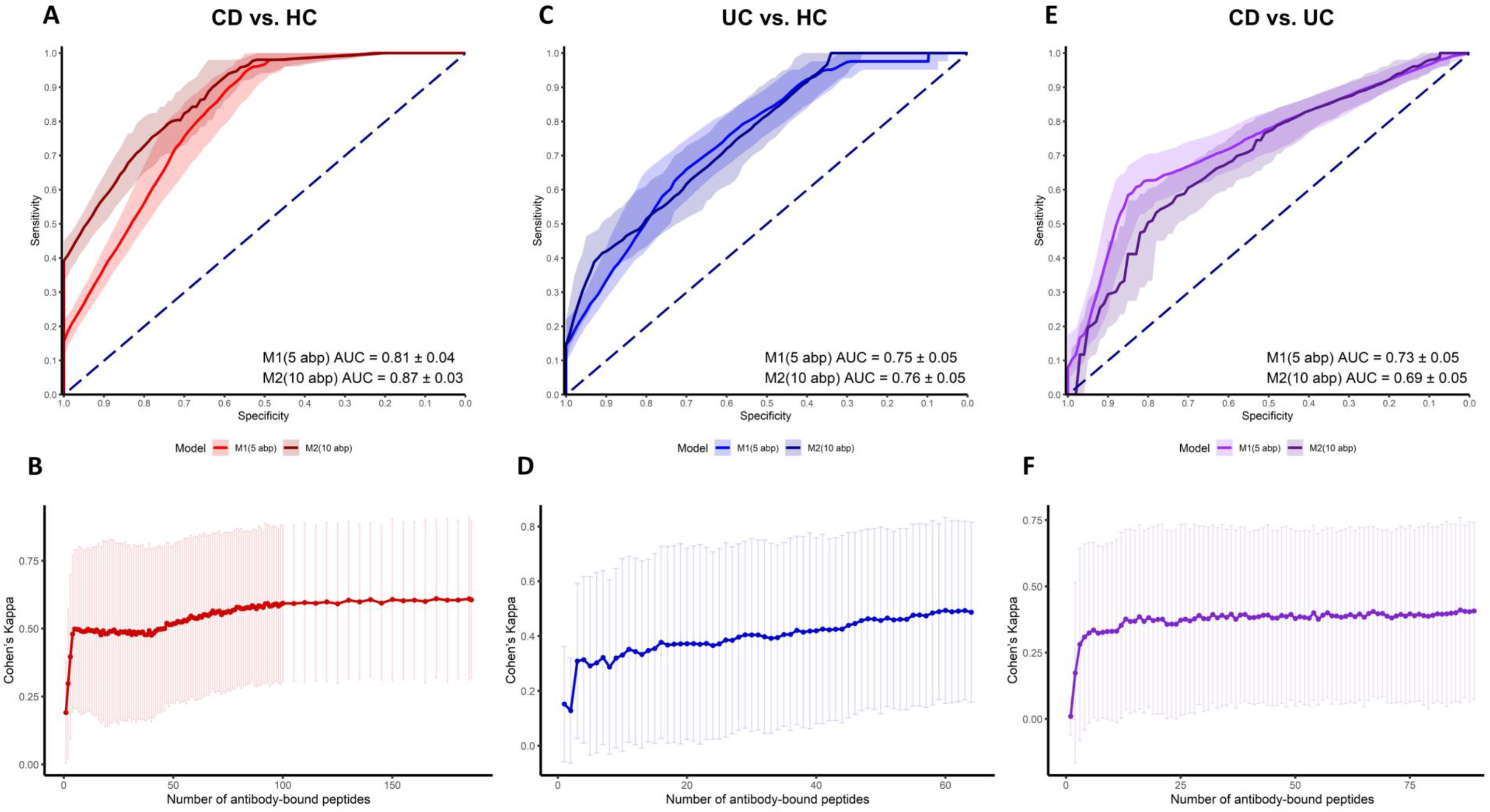
Discriminative capacity of the top five and top ten antibody-bound peptides between patients with CD and healthy controls, patients with UC and healthy controls and patients with CD and UC using recursive feature elimination (RFE). (**A, C, E**) ROC curves demonstrating the discriminative capacity of the top five and top ten antibody-bound peptides in classifying between patients with CD and HC (**A**), patients with UC and HC (**C**) and patients with CD and UC (**E**) in the test set (20% of the data). (**B, D, F**) RFE plots demonstrating the association between the number of antibody-bound peptides entering the classification model and Cohen’s Kappa value, representing a balanced metric between positive and negative predictive values. Abbreviations: ABP, antibody-bound peptides; AUC, area under the curve; CD, Crohn’s disease; HC, (age-and sex-matched) healthy controls; UC, ulcerative colitis.

### Models discriminating patients with CD from healthy controls demonstrate a high degree of external generalizability

To test the external validity of our classification results, we applied our classification models to an external, publicly available dataset from a population-based cohort featuring data on 1,874 PhIP-seq–based antibody-bound peptides from 997 healthy individuals derived from the same microbiota phage library employed in the present study (Vogl et al., 2021). The classification models discriminating patients with CD from healthy controls showed fairly accurate performance on the external dataset (using only the negative side of the classification as there were no patients with IBD in this dataset), whereas this was not observed for the classification models discriminating between patients with UC and healthy controls (**Table S9**). For instance, antibody epitope repertoires demonstrated a highly accurate discrimination between patients with CD and healthy controls from the external dataset (accuracy = 0.82 vs. 0.80 in our own test set), which was not the case for the discrimination between patients with UC and these healthy controls (accuracy = 0.29 vs. 0.70 in our own test set). Similarly, when prioritizing the top-five and top-ten antibody-bound peptides using the RFE-optimized models classifying patients with CD from the external controls, we obtained high accuracies (top five: accuracy = 0.91 vs. 0.72 in our own test set; top ten: accuracy = 0.98 vs. 0.76 in our own test set), which was again not the case when discriminating patients with UC from external controls (top five: accuracy = 0.00 vs. 0.68; top ten: accuracy = 0.12 vs. 0.66) (**Table S11**). Overall, this analysis shows that both the full set and the top-five and top-ten selected antibody-bound peptides are able to accurately classify patients with CD from healthy controls and that this generalizes well to healthy controls from an independent external control cohort.

### Patients with ileal and stricturing CD show distinct antibody responses against specific *Lachnospiraceae* flagellins

Many of the flagellin-derived antigen peptides that were significantly overrepresented in CD (Fig. 3) were particularly enriched in patients with CD with ileal involvement (*n* = 187) as compared to patients with purely colonic CD or UC (*n* = 299). In total, 59 antibody-bound peptides were differentially abundant between patients with ileal CD vs. colonic IBD, of which 95% was overrepresented in ileal CD (FDR < 0.05, Fig. 6A, **Table S14**). Most of these differentially abundant flagellin antigen peptides were derived from bacteria belonging to the order Clostridiales, especially to *Clostridium* clusters XIVa and IV, of which most were in the family Lachnospiraceae, including *Roseburia* spp. (26%), *Lachnospira* spp. (21%), *Clostridium* spp. (19%) and *Eubacterium* spp. (11%), as well as flagellin antigens from *Agathobacter rectalis* and *Butyrivibrio crossotus*. A few significantly enriched flagellin-derived antigen peptides belonged to pathogenic species, including *Legionella pneumophila*, *Borrelia burgdorferi*, *Burkholderia pseudomallei* and *Yersinia enterocolitica*. Furthermore, a few antibody-bound peptides represented ATP-ase and permease components of ATP-binding cassette (ABC)-type transport systems of *Anaerostipes hadrus* and *Coprococcus catus*, both belonging to the family Lachnospiraceae and both well-known butyrate producers. Together, these 59 antibody-bound peptides demonstrated high accuracy in differentiating between patients with ileal CD vs. patients with colonic IBD (AUC = 0.82, Fig. 6A).

**Figure 6.**
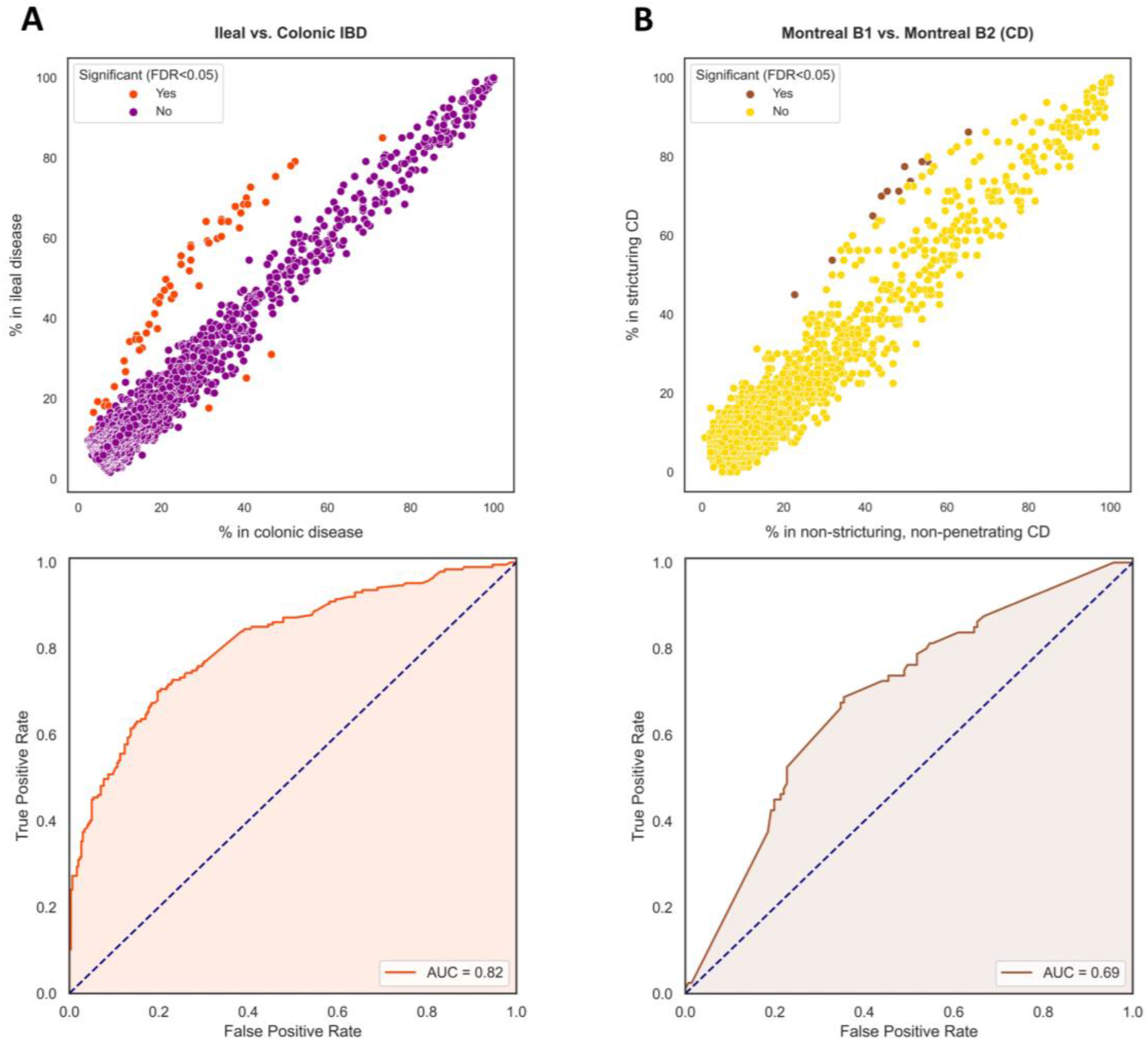
Ileal and stricturing CD disease phenotypes show distinct antibody responses against specific *Lachnospiraceae* flagellins. (**A**) A distinct cluster of antibody-bound peptides (orange-red dots) clearly separates patients with ileal disease involvement (Montreal L1/L3, CD) from patients with IBD with solely colonic disease involvement (Montreal L2 for CD and Montreal E for UC). (**B**) Patients with CD with a stricturing disease phenotype (Montreal B2) demonstrate characteristic antibody responses against a few specific bacterial flagellins. Abbreviations: CD, Crohn’s disease; FDR, false discovery rate; IBD, inflammatory bowel disease; Montreal B1, non-stricturing, non-penetrating CD; Montreal B2, stricturing CD.

In patients with stricturing CD (Montreal classification category B2, *n* = 80), 12 different antibody-bound peptides were significantly enriched compared to patients with non-stricturing, non-penetrating CD (Montreal category B1, *n* = 141) (FDR < 0.05) (Fig. 6B, **Table S15**). All of these peptides originated from bacterial flagellins, especially those from anaerobic, butyrate-producing bacteria, including *Roseburia* spp., *A. rectalis*, *Lachnospira pectinoschiza*, *Eubacterium* spp. and undefined *Lachnospiraceae*. In addition, a few peptides from flagellins of *Clostridium* spp. And *L. pneumophila* were among those differentially enriched in patients with stricturing CD. Together, these 12 antibody-bound peptides were all overrepresented in patients with ileal disease involvement (and were all among the 59 differentially abundant peptides between ileal and colonic IBD, **Table S14**). However, only these 12 differentially abundant antibody-bound peptides demonstrated moderate accuracy to differentiate between patients with stricturing CD from those with non-stricturing, non-penetrating CD (AUC = 0.69, Fig. 6B).

### Surgical history has a modest impact on antibody responses in patients with IBD

Patients with CD who had a history of ileocecal resection (*n* = 64) and patients with IBD who underwent colon resections (*n* = 65) did not demonstrate statistically significant differences in antibody-bound peptides (FDR < 0.05, **Tables S16–17**) compared to patients without surgical history. However, several top-associated antibody-bound peptides (although not passing the FDR threshold) are still noteworthy considering the type of surgical procedure performed (nominal *P*-values < 0.05). For example, patients with CD who underwent an ileocecal resection, more or less reflecting ileal disease involvement, more frequently demonstrated antibody responses against specific flagellin-coated bacteria, including both commensal (including *Lachnospiraceae*, *Roseburia*, *Eubacterium* and *Agathobacter*) and pathogenic (e.g. *Salmonella*, *Yersinia*, *Legionellaceae* and *Helicobacter pylori*) bacterial antigens. In addition, increased antibody responses against specific bacterial virulence factors, including autolysins, bacterial cell wall components and proteins involved in carbohydrate recognition (e.g. peptidoglycan-binding proteins and alpha-glucosidases, respectively). In contrast, peptides representing human rhinoviruses, zinc metalloproteases and several belonging to *Streptococcus pneumoniae* (surface proteins, N-acetylglucosaminidase, autolysins) were less frequently observed in patients with a history of ileocecal resection.

Patients who had undergone colon resections in the past showed consistently less frequent antibody responses against EBV nuclear proteins (mainly Epstein-Barr nuclear antigen 1, EBNA-1) and peptides from the type III secretion system (T3SS) of enteropathogenic *E. coli* (EPEC), which mediate the adherence to and invasion of intestinal epithelial cells. More specifically, the latter peptides belong to the EspD secretor protein and the translocated intimin receptor (Tir) of EPEC serotypes O157:H7 and O127:H6.

Indeed, previous studies have shown that colonies with *E. coli* species belonging to phylogenetic group B2 are commonly isolated from patients with IBD with colonic disease involvement, particularly patients with (left-sided) UC (Kotlowski et al., 2007; Petersen et al., 2009; Petersen et al., 2015).

### Anti-*Saccharomyces cerevisiae* antibodies and established IBD-associated anti-flagellin antibodies associate with the PhIP-seq–based anti-flagellin antibody signature in patients with CD

Patients with CD show consistently higher antibody reactivity against antigens of the yeast *Saccharomyces cerevisiae* (ASCA). In the present cohort, serological ASCA measurements (IgA/IgG, as measured by ELISA) were available for most patients (93%) at time of sampling. Percentages of ASCA positivity in patients with CD were 46% and 43% for IgA and IgG antibodies, respectively, whereas lower prevalence was observed among patients with UC (IgA: 11%; IgG: 17%). ASCA positivity was associated with multiple antibody-bound peptides derived from bacterial flagellins (**Tables S18–19**). In contrast, anti-neutrophil cytoplasmic antibodies (ANCA) (prevalence of titers ≥ 1:80: CD, 10%; UC, 33%) showed no strong signals when related to the antibody-bound peptides (**Table S20**).

Subsequently, we aimed to identify four additional antimicrobial antibodies (anti-*Escherichia coli* outer membrane porin C [anti-OmpC] and the anti-flagellin antibodies anti-CBir1, anti-Fla2 and anti-A4-FlaX) that are characteristic for IBD and commercially available as a serological marker panel (Prometheus^®^ ELISA test, together with ASCA and ANCA). Sequences of anti-A4-Fla2, anti-CBir1, anti-Fla-X and anti-OmpC antibodies showed overlap with 301, 304, 302 and 166 antibody-bound peptide sequences, respectively (e-values < 0.05, **Table S21**). Most of the peptides that showed high sequence homology with anti-A4-Fla2, anti-CBir1 and anti-Fla-X sequences were overrepresented in patients with CD, and all three were very closely related to the same antibody-bound peptides (292 [∼97%] shared antibody-bound peptides) (Suppl. Fig. S2). Among antibody-bound peptides matched with anti-A4-Fla2, anti-CBir-1 and anti-Fla-X antibodies, 71 (∼23%) were among those peptides that were differentially abundant between patients with CD and population controls (**Table S2**, Fig. 3). Notably, antibody-bound peptides that demonstrated a high degree of sequence identity and/or alignment length also represented the strongest differentially abundant peptides between patients with CD and population controls (Suppl. Fig. S3). In contrast, anti-OmpC antibodies showed only a few unique matches with antibody-bound peptides (166 matches with e-values < 0.05), and these peptides also demonstrated very low presences in both patients with IBD and population controls (most < 2%).

### Concordance between antibody responses and microbiome composition

Next, we aimed to investigate the concordance between patients’ serum antibody responses and their fecal microbial composition (Suppl. Figs. S4-S5). We leveraged fecal metagenomics data available for a subset of the present cohort and generated within one year of sampling (*n* = 137). Relative abundances of bacterial taxa were compared between carriers and non-carriers of enriched antibody-bound peptides (**Table S22**), but no significant associations between bacterial taxa and antibodies were observed (FDR-correction for 448,987 tests). However, when evaluating the top-associated microbial taxa (based on nominal *P*-values), the observed associations were considerably smaller in effect size and demonstrated a high degree of discrepancy (i.e. many microbial taxa were not matched to their associated antibody-bound peptides). To further evaluate the impact of microbiome composition on antibody responses, we analyzed the extent to which microbial taxa could explain the variation in antibody responses (**Table S23**). In addition, we performed Principal coordinate analysis (PCoA) on the metagenomics data and defined dysbiosis scores, which represent the median Bray-Curtis dissimilarity distances to a set of reference controls. The first two principal coordinates (PCos) explained a total of 66.6% of the variation in metagenomics data (PCo1: 53.8%; PCo2: 12.8%) and were associated with the presence of IBD (Suppl. Fig. S5A). Patients with IBD showed considerably higher dysbiosis scores compared to healthy individuals, whereas there were no notable differences between IBD subtypes (Suppl. Fig. S5B–C). Importantly, however, dysbiosis scores were not associated with the first ten PCs describing variation in antibody data (PC1 and PC2 shown in Suppl. Fig. 4A). In addition, no differentially abundant antibody-bound peptides were observed when comparing patients with a dysbiotic vs. a non-dysbiotic microbial composition (**Table S24**). Finally, dysbiosis scores were not associated with antibody diversity (i.e. the number of different enriched antibody-bound peptides per patient, Suppl. Fig. S5B).

## Discussion

In this study, we demonstrate that patients with IBD exhibit distinct antibody epitope repertoires compared to individuals derived from the general population. In particular, patients with CD show strong and diverse antibody responses against flagellin-coated bacteria, which were dominated by bacteria belonging to the Lachnospiraceae family. This anti-flagellin antibody signature was particularly present in patients with ileal disease involvement and more complicated disease (e.g. fibrostenotic disease or surgical history). Consequently, antibody epitope repertoires were able to accurately discriminate patients with CD from healthy controls with high discriminative performance, which was not clearly observed for patients with UC. In addition, we examined the concordance between antibody epitope repertoires and gut microbiota composition, but saw no clear associations with individual microbial taxa or microbial dysbiosis. Collectively, our study presents a large-scale, comprehensive characterization of antibody epitope repertoires of patients with IBD, showing a multitude of unique associations with disease phenotypes and representing a valuable information resource to query for systemic immune-based biomarkers for IBD.

Patients with IBD demonstrated selective patterns of antimicrobial antibody reactivity, particularly responses in patients with CD directed against microbiota flagellins. Flagellins are dominant antigens and elicit strong IgG immune responses in CD (Lodes et al., 2004; Christmann et al., 2015). However, the exact nature of the antigens to which these responses are directed to is now gradually being uncovered, although not yet so comprehensively within a large cohort of patients with IBD (Angkeow et al., 2021). More specifically, we found many antibody responses against Lachnospiraceae flagellins (e.g. flagellins of *Roseburia*, *Clostridium*, *Eubacterium* and *Agathobacter* species), particularly in patients with (ileal) CD. These findings corroborate those from previous studies that profiled adaptive immune responses in IBD (Alexander et al., 2021). Many Lachnospiraceae bacteria (belonging to the order Clostridiales, phylum Firmicutes) are typically reduced in patients with CD, similar to what has been observed for *Faecalibacterium prausnitzii*, a member of the Ruminococcaceae family in the Clostridiales order that is closely related to Lachnospiraceae (Vich Vila et al., 2018; Sokol et al., 2008). These bacteria are usually strict anaerobes and well-known butyrate producers, and butyrate has anti-inflammatory and barrier-protective effects on the intestinal epithelium (Furusawa et al., 2013). As such, most flagellated Lachnospiraceae members are commensal symbionts, which makes their strong immunogenicity rather unexpected. However, flagellins are highly immunogenic proteins, and bacterial flagellae are able to facilitate transport across the intestinal epithelial mucus barrier. This cross-barrier transport may, in turn, be particularly enhanced when intestinal barrier integrity is disrupted, which may translate into an increased propensity to be exposed to the adaptive immune system (Arnott et al., 2004). In addition, the intestinal distribution pattern of Lachnospiraceae bacteria largely overlaps with the classical ileocolonic disease localization of CD. The specificity of this anti-flagellin antibody signature for CD but not UC emphasizes that the two diseases have a different pathophysiology, which may be explained by CD-specific immunity (e.g. aberrant Th1-driven immunity) or genetic susceptibility (Alexander et al., 2020). In addition, one may envision potential therapeutic modulation of this anti-flagellin antibody response, as such specific anti-flagellin ‘immunotherapy’ has previously been shown to prevent colitis in mice by selective ablation of flagellin-reactive CD4^+^ T-lymphocytes (Zhao et al., 2020). This principle could be followed to design and test flagellin-directed immunotherapy in patients with IBD.

Previous studies have investigated antimicrobial antibodies as serological predictors of disease (Choung et al., 2016; van Schaik et al., 2013; Torres et al., 2020; Lee et al., 2021). Established anti-flagellin antibodies such as anti-CBir1, anti-Fla-X and anti-A4-Fla2 (also derived from *Lachnospiraceae* flagellins) could accurately predict the development of CD years before the actual diagnosis (Torres et al., 2020). A recent study demonstrated that pre-existent anti-flagellin antibody responses were associated with future development of CD, independent of subclinical inflammation, intestinal barrier function and genetic risk (Lee et al., 2021). This leads to speculate that serum antibody responses preceding CD-onset may constitute one of the earliest pathogenic events, as the latter study only indicated partial mediation by preclinical intestinal inflammation. Although our study was different in the sense that we did not examine a pre-disease cohort but rather patients with established IBD, systemic antibody epitope repertoires showed a highly accurate discrimination between CD and population-derived individuals, suggesting that the application of PhIP-seq technology may also be a robust and powerful approach to predict the onset of CD and to improve our understanding of its preclinical pathogenesis. The size of the PhIP-seq library, the possibility of high-throughput measurements and the rational selection of peptide antigens make PhIP-seq an attractive and promising tool to characterize antibody profiles for these purposes. Notably, stratifying patients by antibody epitope repertoires may potentially aid in primary disease prevention, in a similar manner to that performed for other immune-mediated inflammatory diseases (Herold et al., 2019; Al-Laith et al., 2019).

ASCA and ANCA antibodies are well-known serological markers for CD and UC, respectively, and their presence may predict the development of IBD (Israeli et al., 2005; Zholudev et al., 2004). Although their exact clinical utility remains questionable, they are often included in the serum biomarker panels used to predict or detect IBD (Torres et al., 2020; Zholudev et al., 2004; Mitsuyama et al., 2016). We observed that ASCA IgG-and IgA-antibodies, constituting established serological markers for CD, were strongly associated with increased anti-flagellin antibody peptides compared to ASCA-negative patients. However, this contradicts findings from a recent study that did not find associations between anti-flagellin antibodies and ASCA IgG/IgA (Lee et al., 2021). This may imply that anti-bacterial and anti-fungal antibodies reflect similar antibody reactivity or point to overlap in the ability of bacteria and fungi to elicit antibody responses. However, the definitive role of different antimicrobial antibody responses in the pathophysiology of IBD remains to be determined.

In the present study, we observed no evident similarities, but instead rather spurious associations between gut microbial abundances and serum antibody responses. We can think of several plausible explanations for this observation. For example, serum antibody responses could be elicited from other sites of the body (e.g. the oral cavity, small intestine) beyond the colon or represent long-lasting immunity triggered by transiently present microbes. To what extent these lasting immune effects are captured, however, remains unknown. For instance, antimicrobial antibody responses in patients with a history of ileocecal resection corresponded well to typical shifts of bacterial populations that are known to occur following postoperative microbial recolonization in patients with CD (Machiels et al., 2020; Zhuang et al., 2021). Similarly, patients who underwent colon resections in the past showed decreased antibody responses against EBV, whose presence is associated with severe, refractory colitis and a higher colectomy requirement in patients with UC (Hosomi et al., 2018; Pezhouh et al., 2018). These observations underscore that profound changes in antigen exposures may not be directly reflected by antibody epitope repertoires and instead reflect the long-term adaptability of the adaptive immune system (Marchix et al., 2018). Furthermore, the detection limits of both techniques may show discrepancies: some microbes detected by fecal metagenomics may not necessarily elicit strong antibody responses, whereas highly immunogenic microbes may not all be detected by metagenomics. Although we restricted the microbiome concordance analysis to patients who were profiled within one year of sampling (in both directions), there could still be a certain amount of temporal heterogeneity that overshadows potential associations between the two data entities. The fact that we did not observe strong concordance with the gut microbiome corroborates the findings from a previous study that also found few associations, mostly those involving peptides occurring in < 5% of individuals (Vogl et al., 2021). Furthermore, it should be noted that fecal sampling was employed for generation of metagenomics data, whereas locally present (mucosal) microbial communities are different and may have stronger immune reactivity.

Strengths of the present study included the extensive characterization of the study cohort with multiple layers of information, such as the availability of detailed patient and fecal metagenomics data, which enable detailed assessment of specific antibody signatures. Another highlight pertains to the successful application of the PhIP-seq technology in the context of IBD, providing a high-resolution, high-throughput platform to study human antibody responses and enabling the characterization of both current antibody responses and responses provoked upon past antigen exposure. This temporal stability allows us to capture antibody responses that are saved in our immunological memory, adding a unique resource of biological information to existing technologies such as fluorescence-activated cell sorting (FACS), IgA-seq or BugFACS methods (Andreu-Sánchez et al., manuscript in preparation; Palm et al., 2014; Kau et al., 2015). Previously, the unique longitudinal stability of the PhIP-seq technology was shown to surpass that of metagenomics shotgun sequencing (Vogl et al., 2021). Furthermore, PhIP-seq accurately informs us about the exact nature of the bound antigens, which remain largely unknown using conventional IgA-seq or BugFACS techniques as they usually only provide information about the presence of certain antibody-coated microbes, while capturing only a fraction of the full antibody repertoire. Some limitations of this study also warrant recognition. For example, the majority of patients with IBD in our cohort were in disease remission, which limited our ability to study differences in antibody responses with varying degrees of disease activity or in new-onset patients. In addition, our assessment of disease activity was limited to clinical and serological parameters, as endoscopic data or fecal calprotectin levels were not sufficiently available at time of sampling. Finally, some technical aspects of PhIP-seq limit further characterization of antibody responses, including the length restriction of included peptides (up to 64 amino acids for the present library) and the absence of peptides from conformational epitopes (while all linear epitopes were represented). Second, only protein antigens are incorporated in the library, whereas other immunogenic molecules like lipids and glycans are not represented. However, non-protein antigens usually generate T-cell-independent antibody responses, which are characterized by production of antibodies with lower affinity but higher avidity (Bunker et al., 2019). The current characterization of antibody-bound peptides may therefore represent only strong (high-affinity) and specific (low avidity) immunological responses, which may be expected to be more relevant biomarkers than low-affinity, high-avidity antibody responses. In the same line of thought, detected associations with single peptides should be interpreted cautiously, whereas associations covering multiple peptides for the same protein may be more reliable. In addition, cautious interpretation is warranted with regard to all coagulation-associated peptides (e.g. fibrinogen-binding proteins of *S. aureus*) because patient serum was compared to control plasma, resulting in distinctly underrepresented coagulase-or fibrinogen-related peptides in patients vs. controls, which may have nothing to do with IBD.

Our results provide a new line of evidence for the existence of aberrant immune responses against the gut microbiota in the pathogenesis of IBD. We observed an enrichment of antibody-bound peptides targeting flagellated bacteria in the blood of patients with CD, in particular those patients with ileal disease and more aggressive disease behavior. In the context of IBD, the PhIP-seq technology could be a powerful tool to search for systemic immune-based biomarkers and expose novel immunological targets. Furthermore, integration of PhIP-seq data with other layers of biological information could improve our molecular understanding of IBD and expose leads for mechanistic studies to further disentangle immunopathogenesis. Thus, PhIP-seq could become a valuable clinical application, capturing the interface between host adaptive immunity and gut microbiota and creating an “immunological fingerprint” of patients with IBD. Our results comprehensively demonstrate the vastness, diversity and nature of the human serum antibody epitope repertoire in a large cohort of patients with IBD, showing distinct antibody responses relating to a variety of disease phenotypes.

## Acknowledgments

The authors would like to thank all participants of the 1000IBD cohort. The authors would like to thank Kate Mc Intyre (Scientific Editor, Dept. of Genetics, University Medical Center Groningen) for English editing. The Lifelines Biobank initiative has been made possible by a subsidy from the Dutch Ministry of Health, Welfare and Sport; the Dutch Ministry of Economic Affairs; the University Medical Center Groningen (UMCG, the Netherlands); the University of Groningen and the Northern Provinces of the Netherlands. The authors wish to acknowledge the services of the Lifelines Cohort Study, the contributing research centers delivering data to Lifelines and all study participants.

## Authors’ contributions

Conceptualization: AB, SA-S, TV, SL, SK, INK, AW, ES, JF, AZ, RKW. Investigation: AB, SA-S, TV, SH, AVV, SL, AK, JF, AZ, RKW. Methodology: AB, SA-S, TV, SH, AVV, AK, JF, AZ, RKW. Funding acquisition: AZ, JF, RKW, CW. Supervision: JF, AZ, RKW. Writing – original draft: ARB. Writing – review and editing: all authors.

## Materials and Methods

### Lead contact

Further information and requests for resources should be directed to the Lead Contact, professor Rinse K. Weersma

### Material availability

A full list of antibody-bound peptides that were enriched in patients with IBD will be made publicly available upon publication.

### Data and code availability

The datasets used and/or analyzed during the current study are available from the corresponding author upon reasonable request. The data for the Groningen IBD cohort can be requested with the accession number EGAS00001002702. The raw metagenomics sequencing data used for this study are available from the European Genome-phenome Archive data repository: 1000IBD cohort (https://www.ebi.ac.uk/ega/datasets/EGAD00001004194) and LL-DEEP cohort (https://www.ebi.ac.uk/ega/datasets/EGAD00001001991). Due to patient confidentiality, the datasets are available upon reasonable request to the University Medical Center Groningen (UMCG) and Lifelines, respectively. All code used for analyses in this study can be found at the following link: https://github.com/GRONINGEN-MICROBIOME-CENTRE/Phip-Seq_LLD-IBD.

### Study cohorts: 1000IBD and Lifelines-DEEP

Serum samples were collected from 497 patients with an established diagnosis of IBD who were included at the outpatient clinic of the UMCG upon participation in the 1000IBD project: a large, multi-omics IBD cohort based in Groningen, the Netherlands (Imhann et al., 2019). Detailed phenotypic information and multi-omics data have been generated for these patients, who were enrolled in the period 2007– 2019. Phenotypic data that were collected included age, gender, body-mass index (BMI, body weight divided by squared height), disease duration, smoking behavior, the Montreal disease classification, medication usage, history of surgery, extra-intestinal manifestations, clinical disease activity and serum C-reactive protein levels, all of which was assessed at time of sampling. Clinical disease activity was recorded as the Harvey-Bradshaw Index for patients with CD and the Simple Clinical Colitis Activity Index for patients with UC. All participants provided written informed consent prior to sample collection. The study was approved by the Institutional Review Board (IRB) of the UMCG, Groningen, the Netherlands (in Dutch: “Medisch Ethische Toetsingscommissie”, METc; IRB no. 2008/338) and was conducted in accordance with the principles of the Declaration of Helsinki (2013). In addition, plasma samples were collected from an independent population-based cohort (Lifelines-DEEP), which is situated in the northern part of the Netherlands (Tigchelaar et al., 2015). In total, 1,326 participants from this cohort were included, after having excluded participants with known irritable bowel syndrome or IBD. The Lifelines-DEEP cohort study was also approved by the IRB of the UMCG (IRB no. M12.113965) and registered at the Lifelines Research Site in Groningen. A detailed description of this cohort can be found elsewhere (Andreu-Sánchez et al., manuscript in preparation).

### PhIP-seq microbiota antigen library

Detailed information on the design, cloning and content of the PhIP-seq microbiota antigen library (covering 244,000 peptide sequences) can be found in a recently published study (Vogl et al., 2021). This microbiota antigen library was measured in combination with a second antigen library covering an additional 100,000 peptide sequences related to phages, viruses and immune epitopes (Leviatan et al., manuscript in preparation). Figure 1 shows a schematic overview of the PhIP-seq procedure and antigen microbiota phage library content.

#### Antigen processing and microbiota phage library cloning

Antigens incorporated into the microbiota phage library were cut into peptides with a maximum length of 64 amino acids (aa) (for the 100,000 variant library: 54 aa) with 20 aa overlap for adjoining peptides. To abide with original codon usage frequencies, antigen sequences were back-translated to DNA using codon usage of *E. coli* (of highly expressed proteins), where restriction sites for cloning within the coding sequence were excluded. If required, coding was repeated in order to achieve unique barcodes in the coding sequence at the 44/75-nucleotide (nt) at the 3’-terminus of each oligonucleotide. As each barcode is a unique sequence (at Hamming distance three (44-nt) or five (75-nt) from all prior sequences in the library), a single read error while sequencing 44-nt barcodes can be corrected, while two read errors can be corrected while sequencing 75-nt barcodes. In addition, a part of the 5’-terminus was sequenced to exclude presence of multiple inserts by verifying the matching of 3’-and 5’-sequences. Alternate codons were randomly applied next to *E. coli* codon usage to allow discrimination between rather similar antigen sequences. Sequencing barcodes were incorporated into the coding sequence to enable usage of the entire oligonucleotide for encoding an antigen, rendering sequencing of the full coding sequence unnecessary. Antigen sequences shorter than 64 aa were encoded by adding a random sequence after the stop codon. Library amplification was initiated by adding the *Eco*RI and *Hind*III restriction sites, antigen sequences, stop codons and annealing sequences together as a 230nt-oligomer pool for the 244,000 variant library (Agilent Technologies, Santa Clara, CA, USA; see **Supplementary Methods** for primer details) and 200nt oligos for the 100,000 variant library (Twist Bioscience, San Francisco, CA, USA). Cloning of the antigen library into the T7 phages was performed following manufacturer’s instructions (T7Select 10-3 cloning kit, catalog no. 70550-3, Merck-Millipore, Burlington, MA, USA).

#### Composition and design of the phage antigen libraries

Two recently developed phage antigen libraries were used in this study, covering a total of 344,000 antigen peptides (Vogl et al., 2021; Leviatan et al., manuscript in preparation). One library consists primarily of microbiota antigens (covering 244,000 peptide sequences from 28,668 different proteins, of which 27,837 proteins were derived from microbiota antigens (excluding human proteins and controls)), while the other library includes phage, viral and food antigens (covering 100,000 antigen variants) (Leviatan et al., manuscript in preparation). Details on the origin and composition of the first library are described in the **Supplementary Methods**. Briefly, about 60% (147,061 oligos) of the microbiota antigen library was dedicated to bacterial genes and strains (122,551 oligos, ∼50%, and 24,510 oligos, ∼10%, respectively); 25% of the library (61,250 oligos) was represented by pathogenic bacterial species (24,500 oligos, ∼10%), antibody-coated bacterial species (22,050 oligos, ∼9%) and probiotic bacteria (14,700 oligos, ∼6%); 10% (24,164 oligos) consisted of virulence factors derived from the virulence factor database (VFDB) and ∼5% (11,525 oligos) were (biological and technical) controls. Concerning the 100,000 variant library, 40,000 peptides represented phages, 30,000 allergens and 30,000 antigens derived from the Immune Epitope Database (IEDB) (Leviatan et al., manuscript in preparation). In general, antigens were rationally selected by prioritizing previously reported bacterial antigens and strains known to trigger antibody responses and potential uncharacterized antigens, while focusing on antigens having a high chance of being exposed to the host immune system, e.g. antigens from bacteria with high abundances, membrane proteins, secreted proteins and motility proteins. Selection of antigens was performed without thought of sequence length, necessitating splitting into peptides with a given limit for DNA oligonucleotide synthesis of 230 nt.

#### Immunoprecipitation and next-generation sequencing

Experimental procedures for immunoprecipitation (IP) and next-generation sequencing (NGS) were performed as described previously, but with minor modifications (Vogl et al., 2021; Mohan et al., 2018). Polymerase chain reaction (PCR) plates used for bead transfer and washing were blocked with 150 μL bovine serum albumin (BSA, 30 g/L in Dulbecco’s phosphate-buffered saline at 4°C overnight incubation). Next, BSA was added to the diluted phage/antibody mixtures for IP at a concentration of 2 g/L. Phage wash buffer that was used for IP contained 0.1 wt/vol% IPEGAL CA 630 (Sigma-Aldrich, catalog no. 13021). Subsequently, antibodies (3 μg of serum IgG antibodies as measured by ELISA) and phages (4,000-fold phage coverage per library variant) were mixed after optimization procedures that were performed to determine the optimal amounts of antibodies and phages for IP. The microbiota antigen library was mixed with a 200 nt 100,000 variant pool in a 2:1 ratio. 96-well plates were incubated with the phage/antibody mixtures at 4°C while mixed on an overhead rotator. Protein A and G magnetic beads were incubated overnight on a rotator at 4°C, and 40 μL of this mixture was added to the plates in a 1:1 ratio (Thermo-Fisher Scientific, catalog nos. 10008D and 10009D) according to manufacturer’s instructions. Beads were put into PCR plates and washed twice after 4-h incubation using a Tecan freedom Evo liquid-handling robot with filter tips. Pooled Illumina amplicon sequencing was performed with PCR amplifications using Q5 polymerase (New England Biolabs, catalog no. M0493L) (see **Supplementary Methods** for primer details), and PCR products were paired-end sequenced on an Illumina NextSeq machine.

### Metagenomics shotgun sequencing (MGS) of fecal samples

Fecal metagenomics data were available from a subset of the IBD cohort taken within 1 year of serum sampling (*n* = 137) as well as from the LL-DEEP cohort (Andreu-Sánchez et al., manuscript in preparation). Patients and participants were requested to collect and freeze fecal samples at their homes, after which the samples were picked up, transported on dry ice and stored at −80°C until further analysis. Microbial taxonomy was determined with high-resolution whole-genome MGS. DNA extraction procedures and MGS sequencing with the Illumina HiSeq platform were performed as described previously (Vich Vila et al., 2018; Zhernakova et al., 2016). Genomic library preparation was conducted using the Nextera XT Library preparation kit. Removal of adapters and trimming of the ends of metagenomics reads was performed using Trimmomatic (v.0.32). Cleaned metagenomic reads were further processed through a bioinformatic pipeline as described previously (Vich Vila et al., 2018; Bolger et al., 2014). Taxonomic profiling was performed using MetaPhlAn3 with default parameters, and microbial compositions were expressed as relative abundances in the fecal samples (Beghini et al., 2021). Microbial relative abundances were transformed using centered log-ratios (CLR) transformation. Bacteria not present in at least 10% of samples were discarded.

### Statistical analysis

Descriptive data were presented as mean ± standard deviation (SD), median [interquartile range, IQR] or as proportions *n* with corresponding percentages (%). Demographic and clinical characteristics were compared between groups using Mann-Whitney *U*-tests, chi-squared tests or Fisher’s exact tests (if *n* observations were <10). Two-tailed *P*-values ≤ 0.05 were considered statistically significant. PCA was performed for dimensionality reduction to identify distinct clusters, followed by an assessment of their potential determinants. A *k*-means clustering algorithm (*k* = 2) was additionally performed on the dataset (PCs 1 and 2) to label the observed clusters. To assess the associations between patient phenotypic and clinical factors and the presence of antibody-bound peptides, logistic regression analysis was performed while adjusting for the effect of age and sex. Adjustment for multiple comparisons was performed using the Benjamini-Hochberg method (Benjamini and Hochberg, 1995), where associations with FDR < 5% were considered statistically significant. Using GenBank^®^, we queried the corresponding amino acid sequences of four additional IBD-associated antibodies (anti-*Escherichia coli* outer membrane porin C [anti-OmpC] and the anti-flagellin antibodies anti-CBir1, anti-Fla2 and anti-A4-FlaX) and used the *blastp*-application from BLAST^®^ (v.2.10.1) to analyze sequence homology with the peptide amino acid sequences incorporated in the PhIP-seq library. Classification analysis was performed using logistic regression with elastic net penalty (optimizing the *alpha (*elastic net regularization mixing) and *lambda* (regularization strength) parameters) using the *Caret* package (v.6.0-90) in R (v.4.2.1). In addition, three different machine learning algorithms were used to compare classification performance: GBM (package *gbm_2.1.8*, with optimization of number of trees and interaction depth), a SVM model with radial kernel function (package *kernlab* v.0.9-29, with optimization of the cost (C) parameter) and neural networks using model averaging (avNNet, package *nnet* v.7.3-16, with optimization of network nodes and decay value). Input data, consisting of antibody-bound peptides from 256 CD patients, 207 UC patients and equal numbers of age-and sex-matched healthy controls were randomly partitioned into training (80%) and test (20%) sets. The training set was pre-processed by removing antibody-bound peptides that were highly correlated (Pearson’s correlation coefficient ≥ 0.99), showed zero-variance, were present in <1% or >99% of data, or did not show significant difference between classes (Fisher’s exact tests, *P*>0.005). After these pre-processing steps, final input data consisted of 186 antibody-bound peptides for CD vs. healthy controls, 64 for UC vs. healthy controls and 89 for CD vs. UC classification tasks. The training set was used for training the models, including parameter optimization by five repeats of five-fold cross-validation in order to maximize Cohen’s Kappa value (which is a balanced metric of positive and negative predictive values), whereas the test set was solely used for evaluating model performance. Classification performance was assessed by calculating receiver operating characteristics (ROC) statistics and corresponding evaluation metrics, including the area under the curve (AUC), sensitivity, specificity, PPV, NPV, the F1-score (harmonic mean of precision and recall), Cohen’s Kappa and overall classification accuracy. All classification performance metrics were reported for the test set (metrics for training sets after internal cross-validation are provided in the corresponding supplementary tables). Feature selection from the classification models was performed using RFE with resampling in order to identify and quantify the relative importance of antibody-bound peptides that contributed most to the classification as well as to minimize the chance of potential overfitting and maximize performance. Finally, an additional dataset from a previously published population-based cohort study (featuring data on 1,874 antibody-bound peptides from 997 healthy individuals) was used as an external test set to evaluate the generalization of the models to a geographically distinct population (Vogl et al., 2021). Performance of models using the top 5 and top 10 antibody-bound peptides obtained using RFE was compared to the models using all antibody-bound peptides by DeLong’s test, which is a nonparametric test for comparison of AUCs of ROC curves, implemented in the *pROC* package (v.1.18.0) in R.

Statistical analyses were performed using the Python programming language (v.3.8.5, Python Software Foundation, https://www.python.org) using the *pandas* (v.1.2.3), *numpy* (v.1.20.0), *statsmodels* (v.0.12.2), *scipy* (v.1.7.0) and *sklearn* (v.0.24.2) packages and using R (v.4.2.1). Data visualization was performed using the *seaborn* (v.0.11.1) and *matplotlib* (v.3.4.1) packages in Python.

#### Data processing of PhIP-seq

Sequencing reads from NGS after IP were downsized to 1.25 million identifiable reads per sample, capturing reads that were within one error of all possible barcodes for which the paired-end matched the identifiable antibody-bound peptide. A minimum of 750,000 reads was required for data analysis in the case that insufficient reads were obtained. Enrichment of antigens was calculated by comparing the total number of reads per antigen with that of the input read level (when the phage library was sequenced before IP). Each input read level per sample was assumed to generate an output read level null distribution that was fitted using a generalized Poisson distribution. Estimation of its parameters was separately performed for each input read level in each individual sample, and scores were generated while the parameters were fitted to three distribution parameters for all samples, following interpolation for each input read level (Larman et al., 2011). Individual *P*-values were calculated and adjusted for multiple comparisons using Bonferroni correction (adjusted *P ≤* 0.05 considered statistically significant and defined as seropositivity). Fold changes were computed as number of reads for antibody-bound peptides (after IP) *versus* the number of input reads (before IP), which were computed only for significantly enriched antibody-bound peptides (called as seropositive antibodies), whereas the remaining peptides were set to zero. Furthermore, fold changes were only calculated with a minimum of 25 input reads per peptide. Baseline sequencing of the antigen phage library (before IP) was performed at >100-fold coverage. Samples with <200 significantly enriched antigens (compared to input reads) were excluded. Two prevalence filters were applied for selecting antibody-bound peptides to be used in statistical analyses, where we included peptides that appeared in at least 5% but below 95% of subjects in either (IBD or LL-DEEP) cohorts. Finally, to avoid redundancy of antibody-bound peptides with identical sequences, the most prevalent peptide of sequence replicates was chosen, resulting in a final selection of 2,815 antibody-bound peptides for case–control analyses and 2,368 antibody-bound peptides for IBD-cohort-specific analyses.

#### Microbiome association analysis

Microbiome analyses were performed using the *vegan* package in R (v.4.0.3) (Dixon, 2003). The concordance between antibody profiles and fecal microbial composition was determined by comparing the CLR-transformed relative abundances of bacterial taxa between carriers and non-carriers of each antibody-bound peptide using logistic regression analysis, while adjusting for the effects of age and sex. To assess the contributing effect of microbial taxa on the variation in antibody responses, permutational multivariate ANOVA (PERMANOVA) was used while adopting 1,000 permutations (Adonis function). PCoA was performed to characterize the fecal microbial composition and relate this to microbial dysbiosis, which was quantified using a dysbiosis score based on median Bray-Curtis dissimilarities to a reference sample set of metagenomes (Lloyd-Price et al., 2019). Associations between dysbiosis and the presence of antibody-bound peptides were assessed using logistic regression analysis adjusted for age and sex.

## Supplementary Table Legends

**All Supplementary Tables have been uploaded separately for peer-review.**

**Supplementary Table S1.** Associations between the presences of antibody-bound peptides and patient age by comparison of the first (Q1, <28 years) and fourth age quartiles (Q4, >53 years).

**Supplementary Table S2.** Differentially abundant antibody-bound peptides between male (n=166) and female (n=297) patients with IBD.

**Supplementary Table S3.** Differentially abundant antibody-bound peptides between patients with IBD (full cohort) and healthy individuals (logistic regression analysis, adjusted for the effects of age and gender).

**Supplementary Table S4.** Differentially abundant antibody-bound peptides between patients with CD and healthy individuals (logistic regression analysis, adjusted for the effects of age and gender).

**Supplementary Table S5.** Differentially abundant antibody-bound peptides between patients with UC and healthy individuals (logistic regression analysis, adjusted for the effects of age and gender).

**Supplementary Table S6.** Differentially abundant antibody-bound peptides between patients with CD and an equally sized subset of age- and sex-matched healthy controls (logistic regression analysis, adjusted for the effects of age and gender).

**Supplementary Table S7.** Differentially abundant antibody-bound peptides between patients with UC and an equally sized subset of age- and sex-matched healthy controls (logistic regression analysis, adjusted for the effects of age and gender).

**Supplementary Table S8.** Differentially abundant antibody-bound peptides between patients with CD and UC (logistic regression analysis, adjusted for the effects of age and gender).

**Supplementary Table S9.** Summary of evaluation metrics of three different classification models (CD vs HCs, UC vs HCs and CD vs. UC) using four different (machine learning-based) classification methods.

**Supplementary Table S10.** Beta coefficients and relative importances of antibody-bound peptides for all three classification tasks (CD vs HCs, UC vs. HCs and CD vs. UC).

**Supplementary Table S11.** Summary of evaluation metrics of recursive feature elimination (RFE)-optimized classification models (CD vs. HCs, UC vs. HCs and CD vs. UC) when selecting the top five (5) and top ten (10) antibody-bound peptides.

**Supplementary Table S12.** Beta coefficients and relative importances of the top five (5) and top ten (10) antibody-bound peptides contributing to model classification performance obtained using recursive feature elimination (RFE).

**Supplementary Table S13.** Comparison of areas under the curve (AUCs) of the non-optimized logistic regression with elastic net models, the RFE-optimized top 5 antibody-bound peptide-containing model and the RFE-optimized top 10 antibody-bound peptide-containing model (using DeLong’s test).

**Supplementary Table S14.** Differentially abundant antibody-bound peptides between patients with ileal IBD (Montreal L1/L3) and patients with solely colonic IBD (Montreal L2/E).

**Supplementary Table S15.** Differentially abundant antibody-bound peptides between patients with stricturing CD (Montreal B2) and non-stricturing, non-penetrating CD (Montreal B1).

**Supplementary Table S16.** Differentially abundant antibody-bound peptides between patients who had a history of ileocecal resection (ICR) and those who did not have a past ICR.

**Supplementary Table S17.** Differentially abundant antibody-bound peptides between patients who had a history of (partial) colon resections and those who did not have previous colon resections.

**Supplementary Table S18.** Differentially abundant antibody-bound peptides between patients who had IgG antibodies against Saccharomyces cerevisiae (ASCA) at time of sampling.

**Supplementary Table S19.** Differentially abundant antibody-bound peptides between patients who had IgA antibodies against Saccharomyces cerevisiae (ASCA) at time of sampling.

**Supplementary Table S20.** Differentially abundant antibody-bound peptides between patients who had anti-neutrophil citrullinated antibodies (ANCA) at time of sampling.

**Supplementary Table S21.** Analysis of sequence homology between well-established anti-flagellin antibodies (anti-A4-Fla2, anti-Fla-X, anti-CBir1 and anti-OmpC) in IBD and anti-bound peptides as analyzed by PhIP-seq.

**Supplementary Table S22.** Associations between microbial taxa (derived from metagenomics shotgun sequencing [MGS] data) and antibody-bound peptides (logistic regression analysis after partialling out effects of age and gender).

**Supplementary Table S23.** Permutational multivariate analysis of variance (PERMANOVA) or ADONIS analysis demonstrating the explained variance in antibody-bound peptides by different microbial taxa derived from MGS.

**Supplementary Table S24.** Associations between dysbiosis scores (based on Bray-Curtis dissimilarity indices) and the presence of antibody-bound peptides.

## Supplementary Figures

**Supplementary Figure S1.**
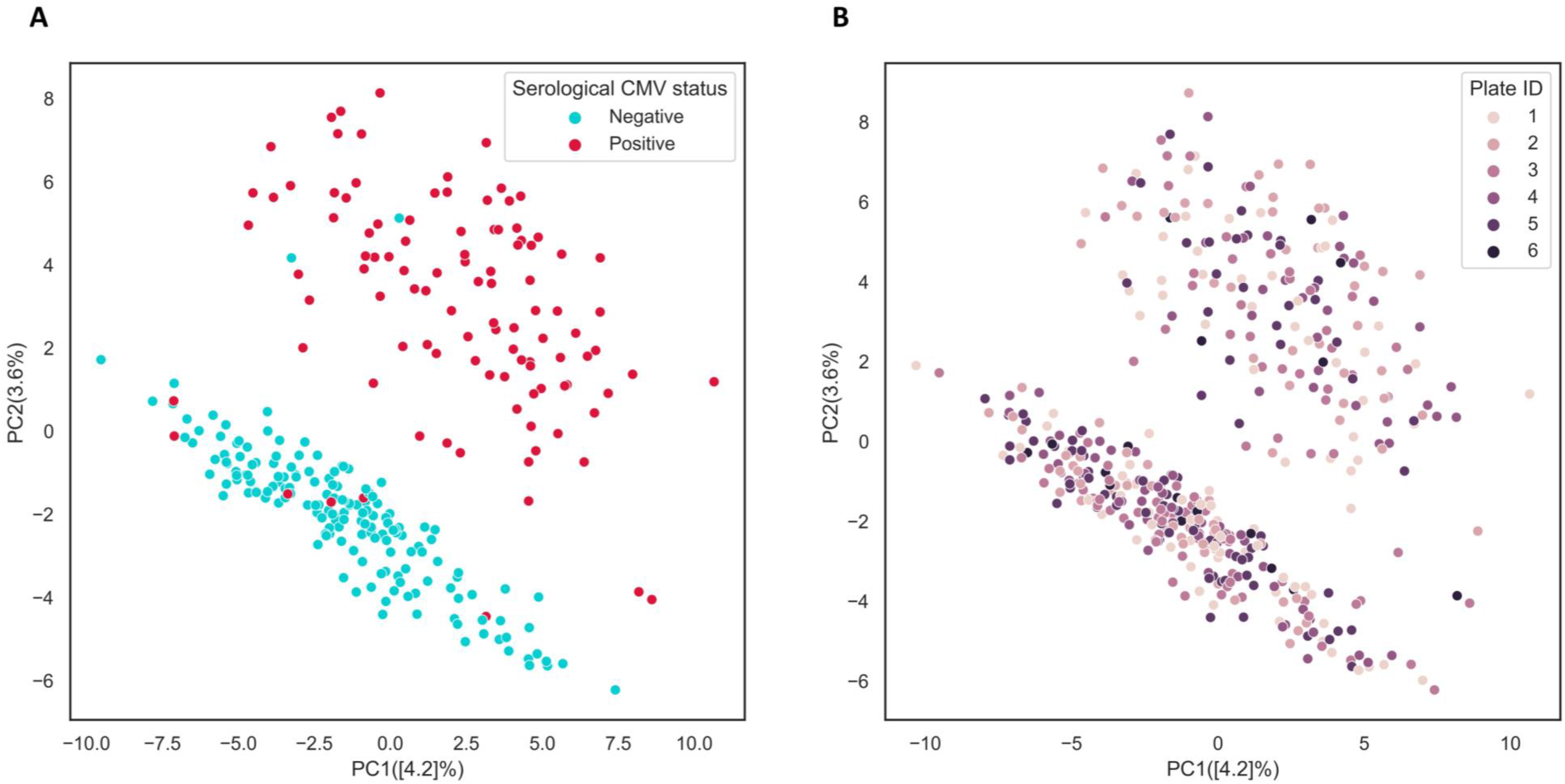
Antibody formation against cytomegalovirus (CMV) determines clustering of the first two principal components (PCs). (**A**) Plate ID, defining the plates on which blood samples were loaded on, does not discriminate between the first two PCs. (**B**) Serological CMV test results available for ∼60% of the IBD cohort, confirmed that antibody responses against CMV antigens determine the clustering of the first two PCs in the PCA of the PhIP-seq data.

**Supplementary Figure S2.**
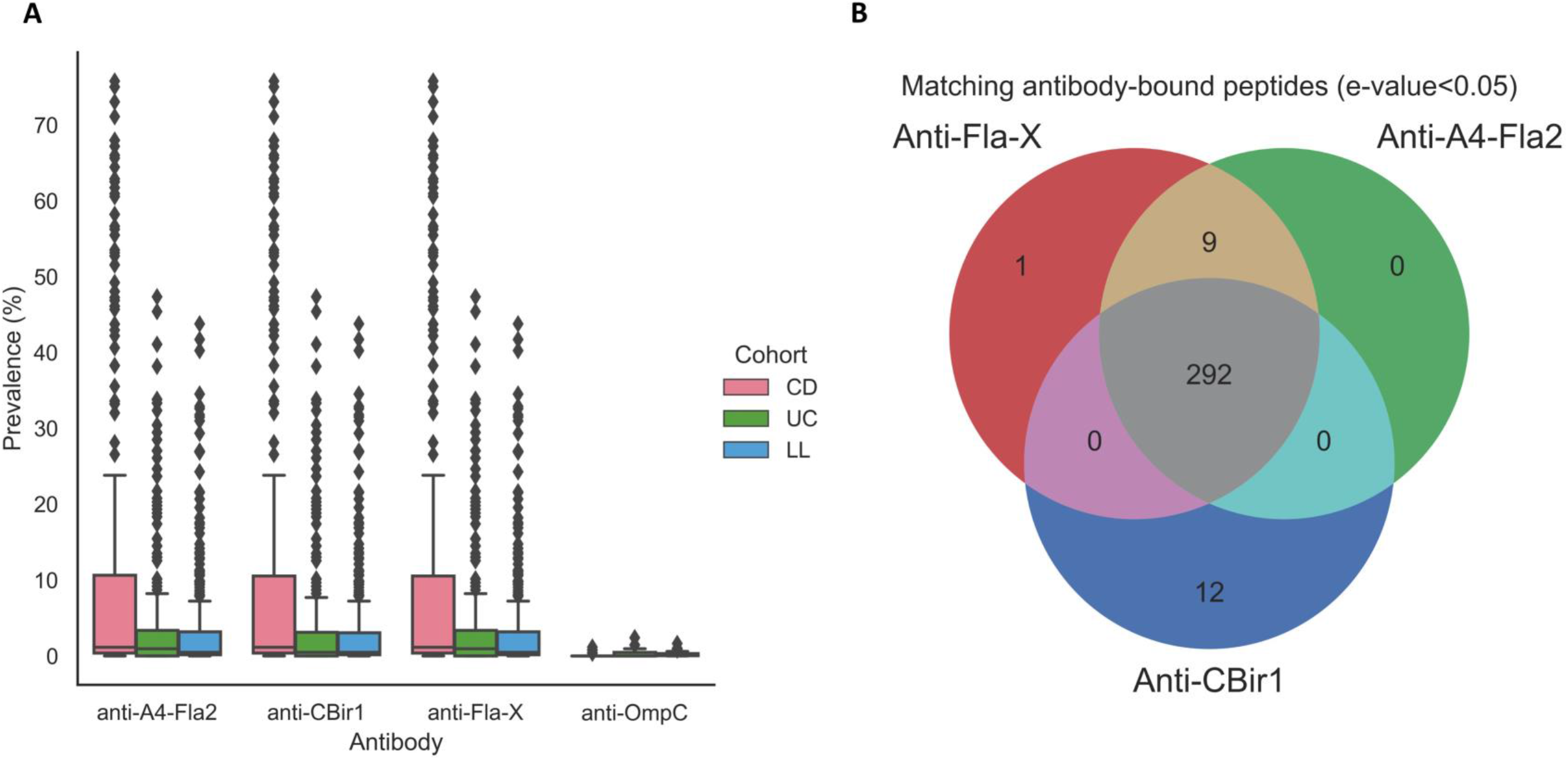
Prevalence of sequence-matched antibody-bound peptides from the PhIP-seq library to the established IBD-associated antimicrobial antibodies anti-A4-Fla2, anti-CBir1, anti-Fla-X and anti-OmpC. **(A)** Patients with CD have higher frequencies of sequence-matched antibody-bound peptides for anti-A4-Fla2, anti-CBir-1 and anti-Fla-X antibodies, whereas matches with anti-OmpC antibodies were rarely present among both patients and population controls. (**B**) Sequence-matched antibody-bound peptides for anti-A4-Fla2, anti-CBir-1 and anti-Fla-X antibodies show a high degree of overlap (∼97%) between all three antibodies.

**Supplementary Figure S3.**
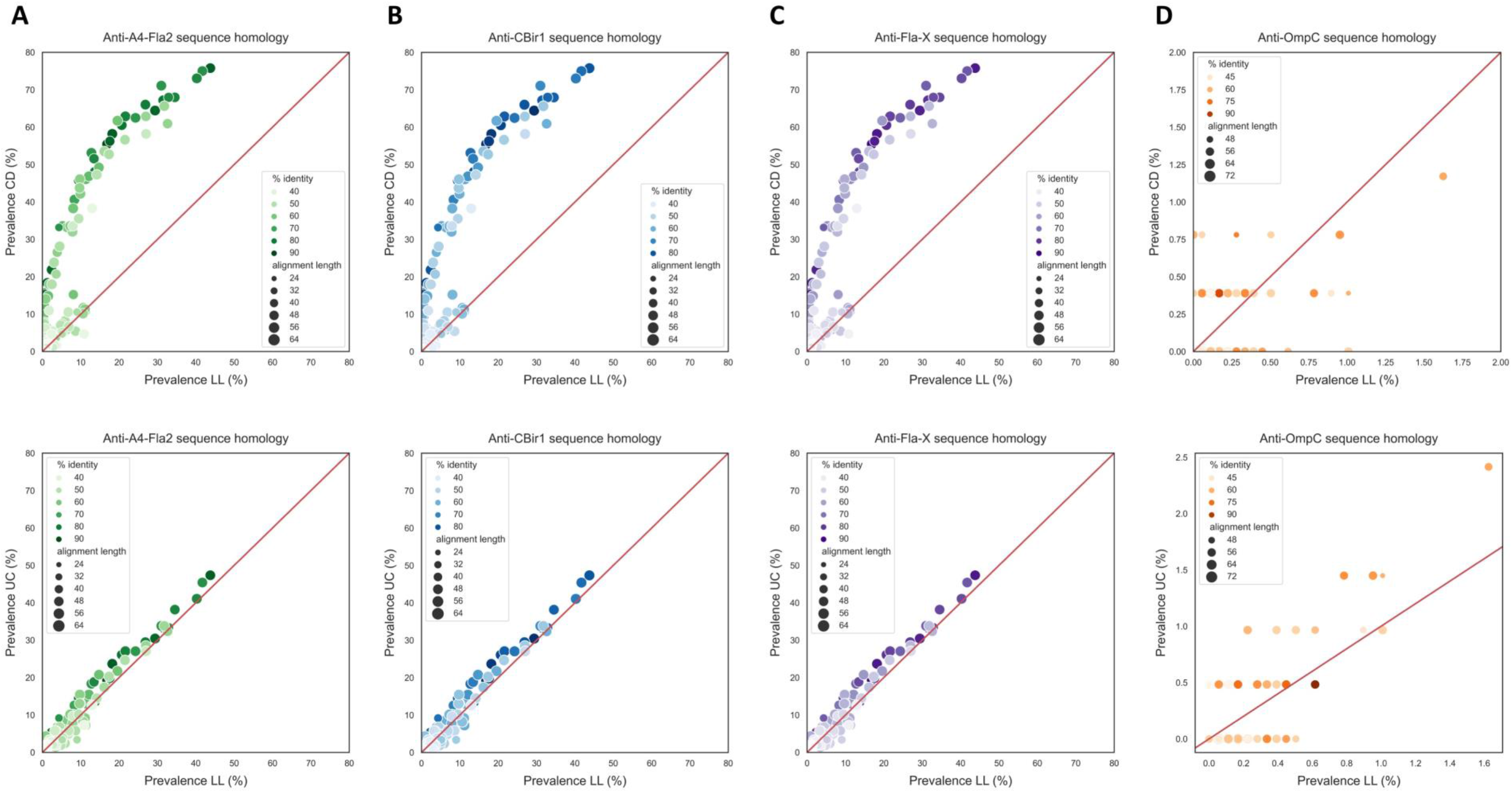
Sequence-matched antibody-bound peptides from the PhIP-seq library to anti-flagellin antibodies anti-A4-Fla2, anti-CBir1 and anti-Fla-X demonstrate a high degree of sequence identity and alignment length, which are among the strongest differentially abundant peptides between patients with CD and population controls. (**A–C**) Percentages of significantly enriched sequence-matched antibody-bound peptides to anti-A4-Fla2 (**A**), anti-CBir-1 (**B**) and anti-Fla-X (**C**) antibodies are considerably higher among patients with CD compared to patients with UC and population controls. (**D**) Anti-OmpC antibodies show only a few unique matches with antibody-bound peptides, which demonstrate very low presence in both patients with IBD and population controls (most <2%).

**Supplementary Figure S4.**
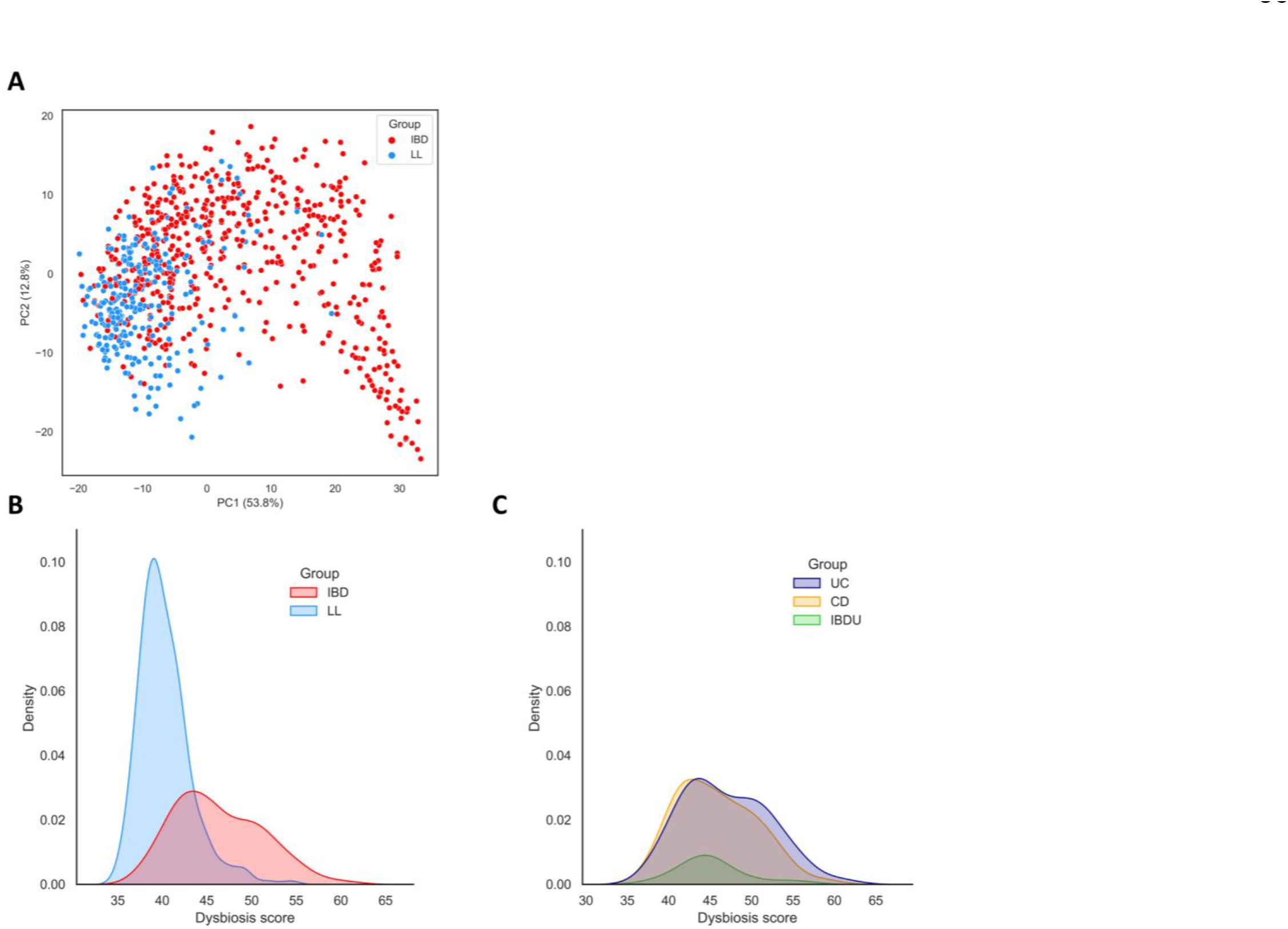
Microbial dysbiosis largely determines variation in fecal metagenomics data and is a hallmark of IBD. (**A**) Principal component analysis (PCA) demonstrating that study cohort (IBD vs. LLD healthy individuals) significantly associates with the first two PCs. (**B–C**) Patients with IBD demonstrate higher dysbiosis scores compared to healthy individuals (**B**), whereas there are no notable differences between IBD subtypes (CD, UC and IBD-U) (**C**).

**Supplementary Figure S5.**
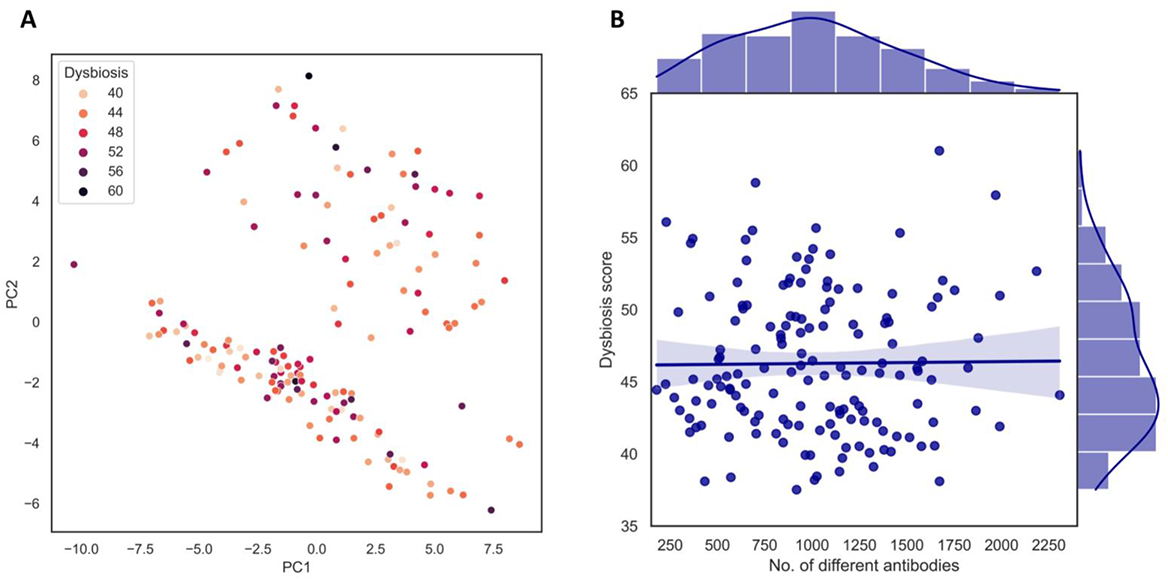
Microbiome dysbiosis is not associated with serum antibody responses in patients with IBD. (**A**) PCA demonstrating that dysbiosis scores do not associate with the first two PCs describing variation in antibody epitope repertoires. (**B**) Microbiome dysbiosis is not associated with diversity in antibody responses (i.e. the number of different enriched antibody-bound peptides per patient).

## Supplementary Methods

### Antigen phage library and Illumina sequencing primers

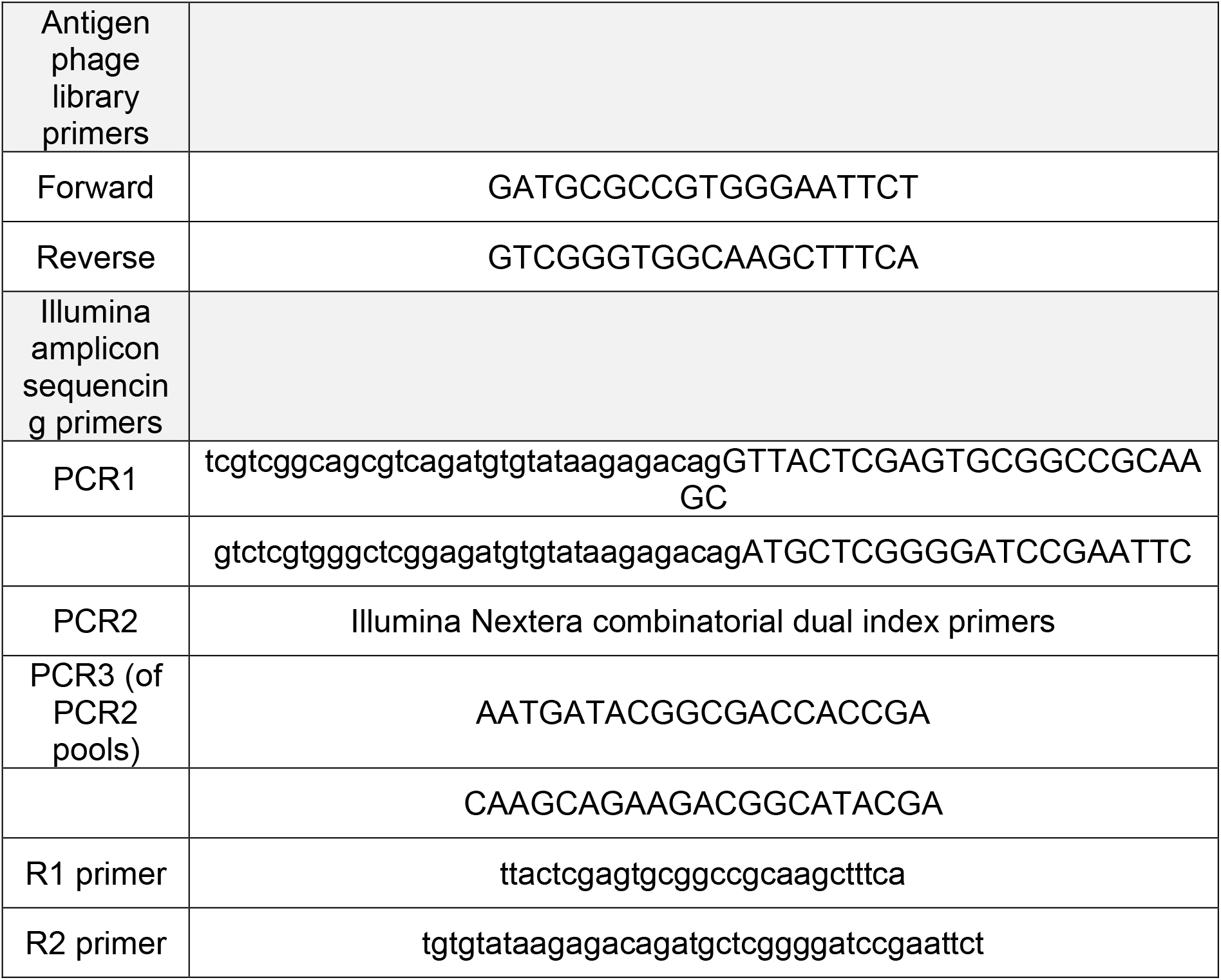

### Origin and composition of the antigen phage microbiota (“Agilent”) library

The composition of the antigen phage microbiota library and of the antigen sequences it contains can be roughly divided into seven different parts, which are summarized below. Full details on the library composition can be found elsewhere (Vogl et al., 2021).

#### (Part I) Bacterial genes (∼50%)

Roughly half of the full antigen phage library is dedicated to antigens from bacterial genes, which were derived from metagenomics shotgun sequencing data of an Israeli population-based cohort, as previously described (Vogl et al., 2021; Zeevi et al., 2015). Mapping of bacterial genes and calculation of their relative abundances was performed using the Integrated Gene Catalog database (Zeevi et al., 2015), and bacterial genes of this background cohort consisted of approximately 4 million different mapped genes. Antigens from these genes were selected based on abundance and annotation, as described previously (Vogl et al., 2021).

#### (Part II) Bacterial strains (∼10%)

Several commonly abundant bacterial strains were added to the library using MetaPhlAn2, a bioinformatics tool, to characterize the taxonomic composition of microbial communities derived from metagenomics shotgun sequencing data (Truong et al., 2015). The ten most-abundant bacterial strains were selected using MetaPhlAn2, from which protein antigen sequences were downloaded from the NCBI.

#### (Part III) Pathogenic bacterial species (∼10%)

The selection of pathogenic bacterial species was based on a previously published study in which the 17 most-prevalent gut pathogens were chosen based on a report of the central laboratories of the Israeli Ministry of Health from 2015. Pathogenic species included were *Salmonella enterica* (subsp. Enterica serovar Enteritidis, Infantis and Muenchen), *Campylobacter coli*, *Campylobacter jejuni* (subsp. jejuni, ATCC 700819), *Vibrio cholerae* (AM-19226), *Vibrio vulnificus*, *Shigella flexneri* (2a str. 301), *Shigella sonnei* (53G), *Listeria monocytogenes* (serotypes 1/2a [str. 08-6569], 1/2b [str. 10-0810], 4b [str. F2365]), *Escherichia coli* (serotype O157:H7, str. Sakai), *Staphylococcus aureus* (subsp. aureus, MRSA252), *Bacillus cereus* (ATCC 14579), *Enterococcus faecalis* (V583) and *Clostridium perfringens* (ATCC 13124).

#### (Part IV) Antibody-coated bacterial species (∼9%)

Antibody-coated bacterial species were selected based on a previous study that investigated IgA-coated microbiota in patients with IBD and healthy individuals (Palm et al., 2014). Antigens were incorporated into the phage library if the IgA coating index of the corresponding bacterial species was >10 and their relative abundances surpassed 10^-6^ in at least three patients with IBD or healthy individuals. Following this, nine bacterial species were selected: five from healthy individuals (*Akkermansia muciniphila* [ATCC BAA-835], *Ruminococcus torques* [ATCC 27756], *Dorea formicigenerans* [ATCC 27755], *Ruminococcus bromii* and *Ruminococcus gnavus* [AGR2154]) and four from patients with IBD (*Blautia producta* [ATCC 27340 = DSM 2950], *Streptococcus lutetiensis* [033], *Bacteroides fragilis* [YCH46], and *Haemophilus parainfluenzae* [T3T1]).

#### (Part V) Probiotic bacterial species (∼6%)

The following probiotic bacterial species were rationally selected based on a recently published review (Lebeer et al., 2018): *Lactobacillus rhamnosus* GG, *Lactobacillus acidophilus* (NCFM), *Lactobacillus plantarum* (WCFS1), *Lactobacillus salivarius* (UCC118), *Lactobacillus reuteri* (DSM 20016), *Escherichia coli* Nissle (1917), *Bifidobacterium breve* (UCC2003), *Bifidobacterium longum* (35624), *Bifidobacterium adolescentis* (ATCC 15703), *Bifidobacterium bifidum* (PRL2010), *Lactobacillus fermentum* (IFO 3956), *Lactobacillus gasseri* (ATCC 33323 = JCM 1131) and *Lactobacillus johnsonii* (NCC 533).

#### (Part VI) Virulence factors (∼10%)

Virulence factor antigen sequences were derived from the virulence factor database (VFDB), which contains antigens from pathogenic bacterial species (Chen et al., 2016). Sequences of 2,624 different genes of ‘set A’ of the VFDB were included, which represents established virulence factors confirmed by previous research.

#### (Part VII) Controls (∼5%)

Both biological and technical controls were included in the antigen phage library. For biological controls, we included B-lymphocyte-derived antigens from proteins of infectious diseases and human auto-antigens extracted from the Immune Epitope Database (IEDB), a public repository covering a wide range of well-established antigens. Antigens from B-cell assays of infectious diseases (excluding parasites) were incorporated, amounting to 290 different proteins represented by 4,250 oligos. Antigens from B-cell assays of human autoimmune diseases were selected as negative controls, which accounted for 430 different proteins represented by 7,700 oligos. Additional negative control antigens that were added to the library (∼300 oligos) included antigens of viral proteins (derived from a previous phage-displayed sequencing study (Xu et al., 2015)) and various peptides that were not expected to trigger antibody responses in our cohorts, including antigens from the Ebola virus, and human antigens such as serum albumin, histone proteins, glycolytic enzymes and ribosomal proteins. Further details and experiments using these controls have previously been described (Vogl et al., 2021). Aside from biological positive and negative controls, technical controls (∼550 oligos) were included in the phage library for experimental validation and contained some random sequences that should not be recognized by antibodies. In addition, codon optimization replicate controls (∼350 oligos) were incorporated to exclude potential bias from different DNA oligonucleotides that might represent the same antigen amino acid sequence. Finally, 50 oligos were included that consisted of very short amino acid sequences (<45 aa) to test for the effects of varying length of the random sequences at the 3’-terminus.

## References

Alexander, K.T., Zhao, Q., Reif, M., Rosenberg, A.F., Mannon, P.J., Duck, L.W., and Elson, C.O. (2021). Human Microbiota Flagellins Drive Adaptive Immune Responses in Crohn’s Disease. Gastroenterology 161, 522–535.e6. 10.1053/j.gastro.2021.03.064.

Al-Laith, M., Jasenecova, M., Abraham, S., Bosworth, A., Bruce, I.N., Buckley, C.D., Ciurtin, C., D’Agostino, M.A., Emery, P., Gaston, H., et al. (2019). Arthritis prevention in the pre-clinical phase of RA with abatacept (the APIPPRA study): a multi-centre, randomised, double-blind, parallel-group, placebo-controlled clinical trial protocol. Trials 20, 429. 10.1186/s13063-019-3403-7.

Andreu-Sánchez, S., Bourgonje, A.R., Vogl, T., Kurilshikov, A., Leviatan, S., Ruiz Moreno, A.J., Hu, S., Sinha, T., Vich Vila, A., Klompus, S., et al. (2021). Genetic, environmental and intrinsic determinants of the human antibody epitope repertoire. Manuscript in preparation.

Angkeow, J.W., Monaco, D.R., Chen, A., Venkataraman, T., Jayaraman, S., Valencia, C., Sie, B.M., Liechti, T., Farhadi, P.N., Funez-dePagnier, G., et al. (2021). Prevalence, persistence, and genetics of antibody responses to protein toxins and virulence factors. bioRxiv, 10.1101/2021.10.01.462481.

Arnott, I.D., Landers, C.J., Nimmo, E.J., Drummond, H.E., Smith, B.K.R., Targan, S.R., and Satsangi, J. (2004). Sero-reactivity to microbial components in Crohn’s disease is associated with disease severity and progression, but not NOD2/CARD15 genotype. Am. J. Gastroenterol. 99, 2376–84. 10.1111/j.1572-0241.2004.40417.x.

Beghini, F., McIver, L.J., Blanco-Míguez, A., Dubois, L., Asnicar, F., Maharjan, S., Mailyan, A., Manghi, P., Scholz, M., Thomas, A.M., et al. (2021). Integrating taxonomic, functional, and strain-level profiling of diverse microbial communities with bioBakery 3. Elife 10, e65088. 10.7554/eLife.65088.

Benjamini, Y., and Hochberg, Y. (1995). Controlling the false discovery rate: a practical and powerful approach to multiple testing. J. R. Stat. Soc. Series B Stat. Methodol. 57, 289–300. 10.1111/j.2517-6161.1995.tb02031.x.

Bischoff, S.C., Barbara, G., Buurman, W., Ockhuizen, T., Schulzke, J.D., Serino, M., Tilg, H., Watson, A., and Wells J.M. (2014). Intestinal permeability—a new target for disease prevention and therapy. BMC Gastroenterol. 14, 189. 10.1186/s12876-014-0189-7.

Bolger, A.M., Lohse, M., and Usadel, B. (2014). Trimmomatic: a flexible trimmer for Illumina sequence data. Bioinformatics 30, 2114–20. 10.1093/bioinformatics/btu170.

Bunker, J.J., Drees, C., Watson, A.R., Plunkett, C.H., Nalger, C.R., Schneewind, O., Eren, A.M., and Bendelac, A. (2019). B cell superantigens in the human intestinal microbiota. Sci. Transl. Med. 11, eaau9356. 10.1126/scitranslmed.aau9356.

Chang, J.T. (2020). Pathophysiology of Inflammatory Bowel Diseases. N. Engl. J. Med. 383, 2652–2664. 10.1056/NEJMra2002697.

Choung, R.S., Princen, F., Stockfisch, T.P., Torres, J., Maue, A.C., Porter, C.K., Leon, F., De Vroey, B., Singh, S., Riddle, M.S., et al. (2016). Serologic microbial associated markers can predict Crohn’s disease behaviour years before disease diagnosis. Aliment. Pharmacol. Ther. 43, 1300–10. 10.1111/apt.13641.

Christmann, B.S., Abrahamsson, T.R., Bernstein, C.N., Duck, L.W., Mannon, P.J., Berg, G., Björkstén, B., Jenmalm, M.C., and Elson, C.O. (2015). Human seroreactivity to gut microbiota antigens. J. Allergy Clin. Immunol. 136, 1378–86.e1-5. 10.1016/j.jaci.2015.03.036.

Dixon, P. (2003). VEGAN, a package of R functions for community ecology. J Veg Sci, 14, 927–930. 10.1111/j.1654-1103.2003.tb02228.x.

Frank, D.N., St Amand, A.L., Feldman, R.A., Boedeker, E.C., Harpaz, N., and Pace, N.R. (2007). Molecular-phylogenetic characterization of microbial community imbalances in human inflammatory bowel diseases. Proc. Natl. Acad. Sci. U. S. A. 104, 13780–5. 10.1073/pnas.0706625104.

Fadlallah, J., Sterlin, D., Fieschi, C., Parizot, C., Dorgham, K., El Kafsi, H., Autaa, G., Ghillani-Dalbin, P., Juste, C., Lepage, P., et al. (2019). Synergistic convergence of microbiota-specific systemic IgG and secretory IgA. J. Allergy Clin. Immunol. 143, 1575–1585.e4. 10.1016/j.jaci.2018.09.036.

Franzosa, E.A., Sirota-Madi, A., Avila-Pacheco, J., Fornelos, N., Haiser, H.J., Reinker, S., Vatanen, T., Hall, A.H., Mallick, H., McIver, L.J., et al. (2019). Gut microbiome structure and metabolic activity in inflammatory bowel disease. Nat. Microbiol. 4, 293–305. 10.1038/s41564-018-0306-4.

Furusawa, Y., Obata, Y., Fukuda, S., Endo, T.A., Nakato, G., Takahashi, D., Nakanishi, Y., Uetake, C., Kato, K., Kato, T., et al. (2013). Commensal microbe-derived butyrate induces the differentiation of colonic regulatory T cells. Nature 504, 446–50. 10.1038/nature12721.

Herold, K.C., Bundy, B.N., Long, S.A., Bluestone, J.A., DiMeglio, L.A., Dufort, M.J., Gitelman, S.E., Gottlieb, P.A., Krischer, J.P., Linsley, P.S., et al. (2019). An Anti-CD3 Antibody, Teplizumab, in Relatives at Risk for Type 1 Diabetes. N. Engl. J. Med. 381, 603–613. 10.1056/NEJMoa1902226.

Hosomi, S., Watanabe, K., Nishida, Y., Yamagami, H., Yukawa, T., Otani, K., Nagami, Y., Tanaka, F., Taira, K., Kamata, N., et al. (2018). Combined Infection of Human Herpes Viruses: A Risk Factor for Subsequent Colectomy in Ulcerative Colitis. Inflamm. Bowel Dis. 24, 1307–1315. 10.1093/ibd/izy005.

Imhann, F., van der Velde, K.J., Barbieri, R., Alberts, R., Voskuil, M.D., Vich Vila, A., Collij, V., Spekhorst, L.M., van der Sloot, K.W.J., Peters, V., et al. (2019). The 1000IBD project: multi-omics data of 1000 inflammatory bowel disease patients; data release 1. BMC Gastroenterol. 19, 5. 10.1186/s12876-018-0917-5.

Israeli, E., Grotto, I., Gilburd, B., Balicer, R.D., Goldin, E., Wiik, A., and Shoenfeld, Y. (2005). Anti-Saccharomyces cerevisiae and antineutrophil cytoplasmic antibodies as predictors of inflammatory bowel disease. Gut 54, 1232–6. 10.1136/gut.2004.060228.

Jostins, L., Ripke, S., Weersma, R.K., Duerr, R.H., McGovern, D.P., Hui, K.Y., Lee, J.C., Schumm, L.P., Sharma, Y., Anderson C.A., et al. (2012). Host-microbe interactions have shaped the genetic architecture of inflammatory bowel disease. Nature 491, 119–24. 10.1038/nature11582.

Kau, A.L., Planer, J.D., Liu, J., Rao, S., Yatsunenko, T., Trehan, I., Manary, M.J., Liu, T.C., Stappenbeck, T.S., Maleta, K.M., et al. (2015). Functional characterization of IgA-targeted bacterial taxa from undernourished Malawian children that produce diet-dependent enteropathy. Sci. Transl. Med. 7, 276ra24. 10.1126/scitranslmed.aaa4877.

Kotlowski, R., Bernstein, C.N., Sepehri, S., and Krause, D.O. (2007). High prevalence of Escherichia coli belonging to the B2+D phylogenetic group in inflammatory bowel disease. Gut 56, 669–75. 10.1136/gut.2006.099796.

Larman, H.B., Zhao, Z., Laserson, U., Li, M.Z., Ciccia, A., Gakidis, M.A., Church, G.M., Kesari, S., Leproust, E.M., Solimini, N.L., et al. (2011). Autoantigen discovery with a synthetic human peptidome. Nat. Biotechnol. 29, 535–41. 10.1038/nbt.1856.

Larman, H.B., Laserson, U., Querol, L., Verhaeghen, K., Solimini, N.L., Xu, G.J., Klarenbeek, P.L., Church, G.M., Hafler, D.A., Plenge, R.M., et al. (2013). PhIP-Seq characterization of autoantibodies from patients with multiple sclerosis, type 1 diabetes and rheumatoid arthritis. J. Autoimmun. 43, 1–9.

Lee, S.H., Turpin, W., Espin-Garcia, O., Raygoza Garay, J.A., Smith, M.I., Leibovitzh, H., Goethel, A., Turner, D., Mack, D., Deslandres, C., et al. (2021). Anti-Microbial Antibody Response is Associated With Future Onset of Crohn’s Disease Independent of Biomarkers of Altered Gut Barrier Function, Subclinical Inflammation, and Genetic Risk. Gastroenterology 161, 1540–1551. 10.1053/j.gastro.2021.07.009.

Leviatan, S., Vogl, T., Klompus, S., Kalka, I.N., Weinberger, A., and Segal, E. (2021). Food proteins elicit distinct systemic antibody responses, that associate with dietary intake in healthy individuals. Manuscript in preparation.

Li, H., Limenitakis, J.P., Greiff, V., Yilmaz, B., Schären, O., Urbaniak, C., Zünd, M., Lawson, M.A.E., Young, I.D., Rupp, S., et al. (2020). Mucosal or systemic microbiota exposures shape the B cell repertoire. Nature 584, 274–8. 10.1038/s41586-020-2564-6.

Lloyd-Price, J., Arze, C., Ananthakrishnan, A.N., Schirmer, M., Avila-Pacheco, J., Poon, T.W., Andrews, E., Ajami, N.J., Bonham, K.S., Brislawn, C.J., et al. (2019). Multi-omics of the gut microbial ecosystem in inflammatory bowel diseases. Nature 569, 655–662. 10.1038/s41586-019-1237-9.

Lodes, M.J., Cong, Y., Elson, C.O., Mohamath, R., Landers, C.J., Targan, S.R., Fort, M., and Hershberg, R.M. (2004). Bacterial flagellin is a dominant antigen in Crohn disease. J. Clin. Invest. 113, 1296–306. 10.1172/JCI20295.

Machiels, K., Pozuelo Del Rio, M., Martinez-De la Torre, A., Xie, Z., Pascal Andreu, V., Sabino, J., Santiago, A., Campos, D., Wolthuis, A., D’Hoore, A., et al. (2020). Early Postoperative Endoscopic Recurrence in Crohn’s Disease Is Characterised by Distinct Microbiota Recolonisation. J. Crohns Colitis 14, 1535–1546. 10.1093/ecco-jcc/jjaa081.

Marchix, J., Goddard, G., and Helmrath, M.A. (2018). Host-Gut Microbiota Crosstalk in Intestinal Adaptation. Cell. Mol. Gastroenterol. Hepatol. 6, 149–162. 10.1016/j.jcmgh.2018.01.024.

von Martels, J.Z.H., Sadaghian Sadabad, M., Bourgonje, A.R., Blokzijl, T., Dijkstra, G., Faber, K.N., and Harmsen, H.J.M. (2017). The role of gut microbiota in health and disease: In vitro modeling of host-microbe interactions at the aerobe-anaerobe interphase of the human gut. Anaerobe 44, 3–12. 10.1016/j.anaerobe.2017.01.001.

Mina, M.J., Kula, T., Leng, Y., Li, M., de Vries, R.D., Knip, M., Siljander, H., Rewers, M., Choy, D.F., Wilson, M.S., et al. (2019). Measles virus infection diminishes preexisting antibodies that offer protection from other pathogens. Science 366, 599–606. 10.1126/science.aay6485.

Mitsuyama, K., Niwa, M., Takedatsu, H., Yamasaki, H., Kuwaki, K., Yoshioka, S., Yamauchi, R., Fukunaga, S., Torimura, T. (2016). Antibody markers in the diagnosis of inflammatory bowel disease. World J. Gastroenterol. 22, 1304–10. 10.3748/wjg.v22.i3.1304.

Mohan, D., Wansley, D.L., Sie, B.M., Noon, M.S., Baer, A.N., Laserson, U., and Larman, H.B. (2018). PhIP-Seq characterization of serum antibodies using oligonucleotide-encoded peptidomes. Nat. Protoc. 13, 1958–78. 10.1038/s41596-018-0025-6.

Palm, N.W., de Zoete, M.R., Cullen, T.W., Barry, N.A., Stefanowski, J., Hao, L., Degnan, P.H., Hu, J., Peter, I., Zhang, W., et al. (2014). Immunoglobulin A coating identifies colitogenic bacteria in inflammatory bowel disease. Cell 158, 1000–1010. 10.1016/j.cell.2014.08.006.

Pasolli, E., Asnicar, F., Manara, S., Zolfo, M., Karcher, N., Armanini, F., Beghini, F., Manghi, P., Tett, A., Ghensi, P., et al. (2019). Extensive Unexplored Human Microbiome Diversity Revealed by Over 150,000 Genomes from Metagenomes Spanning Age, Geography, and Lifestyle. Cell 176, 649–62. 10.1016/j.cell.2019.01.001.

Petersen, A.M., Nielsen, E.M., Litrup, E., Brynskov, J., Mirsepasi, H., and Krogfelt, K.A. (2009). A phylogenetic group of Escherichia coli associated with active left-sided inflammatory bowel disease. BMC Microbiol. 9, 171. 10.1186/1471-2180-9-171.

Petersen, A.M., Halkjær, S.I., and Gluud, L.L. (2015). Intestinal colonization with phylogenetic group B2 Escherichia coli related to inflammatory bowel disease: a systematic review and meta-analysis. Scand. J. Gastroenterol. 50, 1199–207. 10.3109/00365521.2015.1028993.

Pezhouh, M.K., Miller, J.A., Sharma, R., Borzik, D., Eze, O., Waters, K., Westerhoff, M.A., Parian, A.M., Lazarev, M.G., and Voltaggio, L. (2018). Refractory inflammatory bowel disease: is there a role for Epstein-Barr virus? A case-controlled study using highly sensitive Epstein-Barr virus-encoded small RNA1 in situ hybridization. Human Pathol. 82, 187–192. 10.1016/j.humpath.2018.08.001.

Salim, S.Y., and Söderholm, J.D. (2011). Importance of disrupted intestinal barrier in inflammatory bowel diseases. Inflamm. Bowel. Dis. 17, 362–81. 10.1002/ibd.21403.

van Schaik, F.D., Oldenburg, B., Hart, A.R., Siersema, P.D., Lindgren, S., Grip, O., Teucher, B., Kaaks, R., Bergmann, M.M., Boeing, H., et al. (2013). Serological markers predict inflammatory bowel disease years before the diagnosis. Gut 62, 683–8. 10.1136/gutjnl-2012-302717.

Sokol, H., Pigneur, B., Watterlot, L., Lakhdari, O., Bermúdez-Humarán, L.G., Gratadoux, J.J., Blugeon, S., Bridonneau, C., Furet, J.P., Corthier, G., et al. (2008). Faecalibacterium prausnitzii is an anti-inflammatory commensal bacterium identified by gut microbiota analysis of Crohn disease patients. Proc. Natl. Acad. Sci. U. S. A. 105, 16731–6. 10.1073/pnas.0804812105.

de Souza, H.S.P., Fiocchi, C., and Iliopoulos, D. (2017). The IBD interactome: an integrated view of aetiology, pathogenesis and therapy. Nat. Rev. Gastroenterol. Hepatol. 14, 739–749. 10.1038/nrgastro.2017.110.

Sterlin, D., Fadlallah, J., Slack, E., and Gorochov, G. (2020). The antibody/microbiota interface in health and disease. Mucosal Immunol. 13, 3–11. 10.1038/s41385-019-0192-y.

Tigchelaar, E.F., Zhernakova, A., Dekens, J.A., Hermes, G., Baranska, A., Mujagic, Z., Swertz, M.A., Muñoz, A.M., Deelen, P., Cénit, M.C., et al. (2015). Cohort profile: LifeLines DEEP, a prospective, general population cohort study in the northern Netherlands: study design and baseline characteristics. BMJ Open 5, e006772. 10.1136/bmjopen-2014-006772.

Torres, J., Petralia, F., Sato, T., Wang, P., Telesco, S.E., Choung, R.S., Strauss, R., Li, X.J., Laird, R.M., Gutierrez, R.L., et al. (2020). Serum Biomarkers Identify Patients Who Will Develop Inflammatory Bowel Diseases Up to 5 Years Before Diagnosis. Gastroenterology 159, 96–104. 10.1053/j.gastro.2020.03.007.

Vich Vila, A., Imhann, F., Collij, V., Jankipersadsing, S.A., Gurry, T., Mujagic, Z., Kurilshikov, A., Bonder, M.J., Jiang, X., Tigchelaar, E.F., et al. (2018) Gut microbiota composition and functional changes in inflammatory bowel disease and irritable bowel syndrome. Sci. Transl. Med. 10, eaap8914. 10.1126/scitranslmed.aap8914.

Vogl, T., Klompus, S., Leviatan, S., Kalka, I.N., Weinberger, A., Wijmenga, C., Fu, J., Zhernakova, A., Weersma, R.K., and Segal, E. (2021). Population-wide diversity and stability of serum antibody epitope repertoires against human microbiota. Nat. Med. 27, 1442–50. 10.1038/s41591-021-01409-3.

Xu, G.J., Kula, T., Xu, Q., Li, M.Z., Vernon, S.D., Ndung’u, T., Ruxrungtham, K., Sanchez, J., Brander, C., Chung, R.T., et al. (2015). Viral immunology. Comprehensive serological profiling of human populations using a synthetic human virome. Science 348, aaa0698. 10.1126/science.aaa0698.

Zeevi, D., Korem, T., Zmora, N., Israeli, D., Rothschild, D., Weinberger, A., Ben-Yacov, O., Lador, D., Avnit-Sagi, T., Lotan-Pompan, M., et al. (2015). Personalized Nutrition by Prediction of Glycemic Responses. Cell 163, 1079–94. 10.1016/j.cell.2015.11.001.

Zhao, Q., Duck, L.W., Huang, F., Alexander, K.L., Maynard, C.L., Mannon, P.J., and Elson, C.O. (2020). CD4+ T cell activation and concomitant mTOR metabolic inhibition can ablate microbiota-specific memory cells and prevent colitis. Sci. Immunol. 5, eabc6373. 10.1126/sciimmunol.abc6373.

Zhernakova, A., Kurilshikov, A., Bonder, M.J., Tigchelaar, E.F., Schirmer, M., Vatanen, T., Mujagic, Z., Vich Vila, A., Falony, G., Vieira-Silva, S., et al. (2016). Population-based metagenomics analysis reveals markers for gut microbiome composition and diversity. Science 352, 565–9. 10.1126/science.aad3369.

Zholudev, A., Zurakowski, D., Young, W., Leichtner, A., and Bousvaros, A. (2004). Serologic testing with ANCA, ASCA, and anti-OmpC in children and young adults with Crohn’s disease and ulcerative colitis: diagnostic value and correlation with disease phenotype. Am. J. Gastroenterol. 99, 2235–41. 10.1111/j.1572-0241.2004.40369.x.

Zhuang, X., Tian, Z., Li, N., Mao, R., Li, X., Zhao, M., Xiong, S., Zeng, Z., Feng, R., and Chen, M. (2021). Gut Microbiota Profiles and Microbial-Based Therapies in Post-operative Crohn’s Disease: A Systematic Review. Front. Med. (Lausanne) 7, 615858. 10.3389/fmed.2020.615858.

## Supplementary References

Chen L, Zheng D, Liu B, Yang J, Jin Q. VFDB 2016: hierarchical and refined dataset for big data analysis—10 years on. Nucleic Acids Res 2016;44(D1):D694–7.

Lebeer S, Bron PA, Marco ML, Van Pijkeren JP, O’Connell Motherway M, Hill C, et al. Identification of probiotic effector molecules: present state and future perspectives. Curr Opin Biotechnol 2018;49:217–223.

Palm NW, de Zoete MR, Cullen TW, Barry NA, Stefanowski J, Hao L, et al. Immunoglobulin A coating identifies colitogenic bacteria in inflammatory bowel disease. Cell 2014;158(5):1000–1010.

Truong DT, Franzosa EA, Tickle TL, Scholz M, Weingart G, Pasolli E, et al. MetaPhlAn2 for enhanced metagenomic taxonomic profiling. Nat Methods 2015;12(10):902–3.

